# Cell-type specialization in the brain is encoded by specific long-range chromatin topologies

**DOI:** 10.1101/2020.04.02.020990

**Authors:** Warren Winick-Ng, Alexander Kukalev, Izabela Harabula, Luna Zea Redondo, Dominik Szabo, Mandy Meijer, Leonid Serebreni, Yingnan Zhang, Simona Bianco, Andrea M. Chiariello, Ibai Irastorza-Azcarate, Luca Fiorillo, Francesco Musella, Christoph J. Thieme, Ehsan Irani, Elena Torlai Triglia, Aleksandra A. Kolodziejczyk, Andreas Abentung, Galina Apostolova, Eleanor J. Paul, Vedran Franke, Rieke Kempfer, Altuna Akalin, Sarah A. Teichmann, Georg Dechant, Mark A. Ungless, Mario Nicodemi, Lonnie Welch, Gonçalo Castelo-Branco, Ana Pombo

## Abstract

Neurons and oligodendrocytes are terminally differentiated cells that sustain cascades of gene activation and repression to execute highly specialized functions, while retaining homeostatic control. To study long-range chromatin folding without disturbing the native tissue environment, we developed Genome Architecture Mapping in combination with immunoselection (immunoGAM), and applied it to three cell types from the adult murine brain: dopaminergic neurons (DNs) from the midbrain, pyramidal glutamatergic neurons (PGNs) from the hippocampus, and oligodendroglia (OLGs) from the cortex. We find cell-type specific 3D chromatin structures that relate with patterns of gene expression at multiple genomic scales, including extensive reorganization of topological domains (TADs) and chromatin compartments. We discover the loss of TAD insulation, or ‘TAD melting’, at long genes (>400 kb) when they are highly transcribed. We find many neuron-specific contacts which contain accessible chromatin regions enriched for putative binding sites for multiple neuronal transcription factors, and which connect cell-type specific genes that are associated with neurodegenerative disorders such as Parkinson’s disease, or specialized functions such as synaptic plasticity and memory. Lastly, sensory receptor genes exhibit increased membership in heterochromatic compartments that establish strong contacts in brain cells. However, their silencing is compromised in a subpopulation of PGNs with molecular signatures of long-term potentiation. Overall, our work shows that the 3D organization of the genome is highly cell-type specific, and essential to better understand mechanisms of gene regulation in highly specialized tissues such as the brain.

## Introduction

The study of three-dimensional (3D) chromatin organization has revealed its intrinsic association with gene regulation and cell function^1^. Genome-wide sequencing approaches such as Hi-C^2^, GAM^3^ and SPRITE^4^ have shown that the genome is organized hierarchically, from specific promoter-enhancer contacts, to topologically associating domains (TADs), to broader compartments of open and closed chromatin (compartments A and B, respectively), up to whole chromosome territories^2–5^. Dynamic changes of chromatin organization during neuronal development have been reported using *in-vitro* differentiated neurons, or glutamatergic neurons obtained after cortical tissue dissociation and FACS isolation from pools of animals, or from mixed cell populations from whole hippocampi^6–8^. However, it has been difficult to assess chromatin structure of specific cell types from the brain, especially without disrupting tissue organization, and in single animals. Understanding 3D chromatin structure of specialized cell populations is of exceptional importance in the brain, where connecting disease-associated genetic variants in non-coding genomic regions with their target genes remains challenging, and where cell state and localization within the tissue can influence both transcriptional and physiological outcomes.

### ImmunoGAM maps 3D genome architecture in specific mouse brain

Here we developed immunoGAM, an extension of the Genome Architecture Mapping (GAM) technology^3^ to perform genome-wide mapping of chromatin topology in specific cell populations. GAM is a ligation-free technology that maps 3D genome topology by extracting and sequencing the genomic DNA content from nuclear cryosections, followed by detection of 3D chromatin interactions from the increased probability of co-segregation of genomic loci across a collection of nuclear slices, each from a single nucleus. GAM was previously applied to mouse embryonic stem cells (mESCs) and shown to capture TADs, A/B compartments and pair-wise contacts across long genomic distances^3^. Recently, we developed a streamlined version of GAM, multiplexGAM, which combines three nuclear slices in each GAM sample^9^. With immunoGAM, we now considerably extend the scope of GAM to directly work in intact tissue without prior dissociation, and with small cell numbers (∼1000 cells)^3, 10^.

To explore how genome folding is related with cell specialization, we applied immunoGAM in three specific brain cell types with diverse functions (**Fig. 1a**). We selected oligodendroglia (oligodendrocytes and their precursors; OLGs) from the somatosensory cortex, as they are involved in myelination of neurons and influence neuronal circuit formation^11^. Pyramidal glutamatergic neurons (PGNs) were selected from the *cornu ammonis 1* (CA1) region of the dorsal hippocampus. PGNs have specialized roles in temporal and spatial memory formation and consolidation, and are activated intrinsically and frequently^12, 13^. We also selected dopaminergic neurons (DNs) from the ventral tegmental area of the midbrain (VTA), which are normally quiescent but are activated during cue-guided reward-based learning^14^. Finally, we compared GAM data from the selected brain cell types to publicly available multiplexGAM data from mESCs^9^ (see **Methods, Supplemental Table 1**).

**Figure 1.**
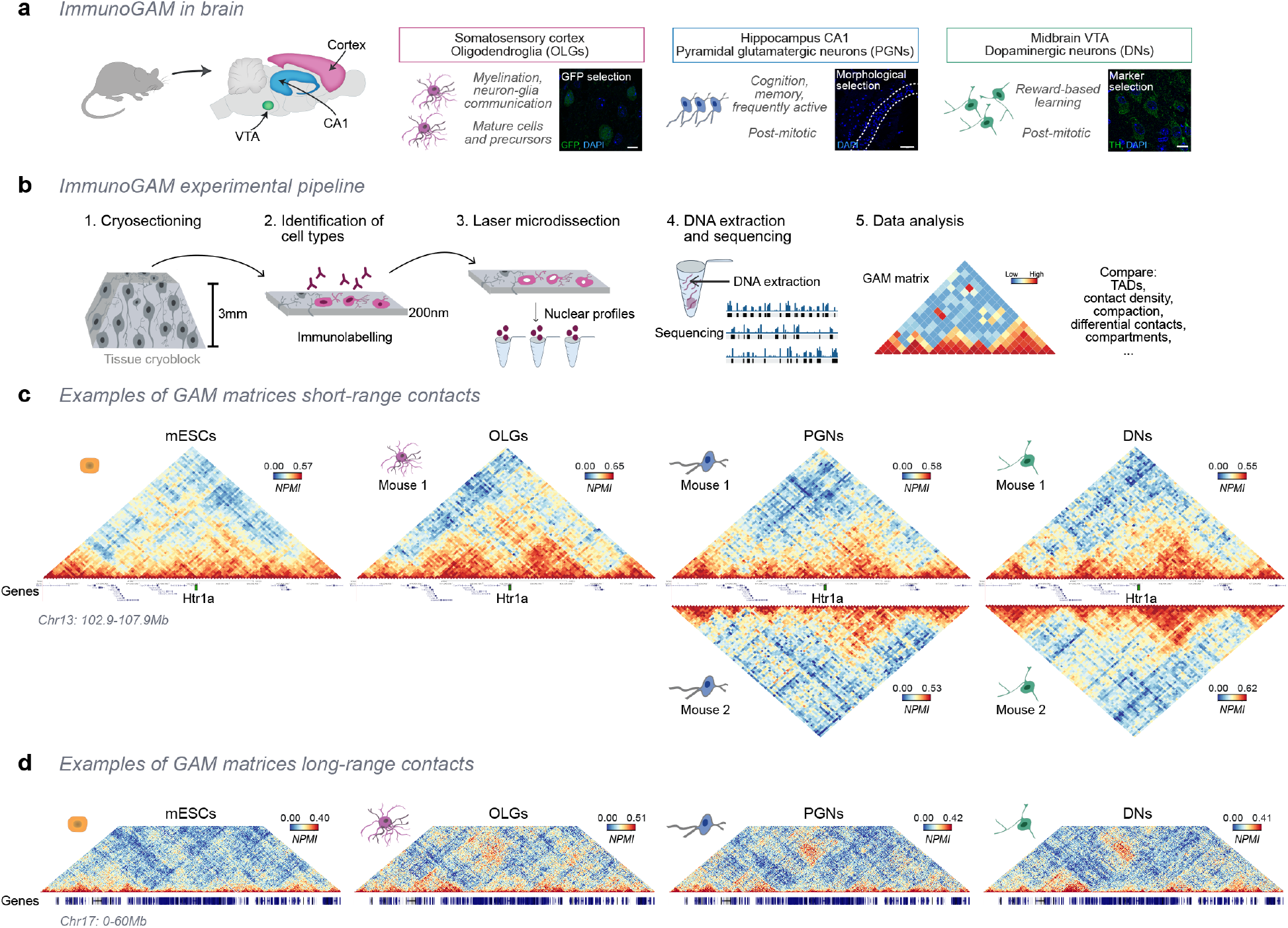
ImmunoGAM captures cell-type specific chromatin contacts in the mouse brain. **a,** Immuno-GAM was applied to oligodendrocytes (OLGs), pyramidal glutamatergic neurons (PGNs), and dopaminergic neurons (DNs) (scale: 10, 100 and 10 µm, respectively). OLGs were selected based on GFP expression, PGNs using tissue morphology (dotted line), and DNs by TH expression. **b**, The immunoGAM experimental workflow. Upon transcardial perfusion with fixative, brain tissues are dissected, cryoprotected, and ultrathin cryosectioned (1). After immunodetection using cell-type specific markers (2), single nuclear profiles are laser microdissected (3), each from a single cell, and collected in groups of three, as described for multiplex-GAM^9^. After extraction of genomic DNA and sequencing (4), the frequency of co-segregation of pairs of genomic loci is used to calculate matrices of pairwise contacts, and to compare the organization of topological domains, chromatin compaction, differential contacts, and compartments (5). **c**, Examples of contact matrices showing cell-type specific differences in local contacts within a 5-Mb genomic region centered on the *Htr1a* gene, which encodes 5-hydroxytryptamine (serotonin) receptor 1A, in mESCs, OLGs, PGNs and DNs. GAM matrices represent co- segregation frequencies of pairs of 50-kb genomic windows using normalized pointwise mutual information (NPMI). Contact maps show reproducible chromatin folding in replicate data collected from single animals for PGNs and DNs. **d**, Exemplar of cell-type specific differences in long-range contacts within a 60-Mb genomic region in chromosome 17. Patches of strong contacts involving two large genomic regions separated by ∼35 Mb are observed in brain cells, but not in mESCs.

Tissues were collected from fixative-perfused animals, cryoprotected by sucrose embedding, and then frozen in liquid nitrogen (**Fig. 1b**). After cryosectioning into ∼230 nm thin tissue slices, the brain cells of interest were immunostained with specific antibodies. For each GAM sample from single animals, we collected nuclear slices from single cells, by laser microdissection. Next, the DNA content of each sample (containing three nuclear slices) was extracted, amplified and sequenced (627-1755 cells passed QC; **Table 1**). A detailed flowchart of the data collection and quality control measures are presented in **Extended Data Fig. 1a-c, and Supplemental Table 2**. We selected a working resolution of 50 kb, based on good sampling of pair-wise co-segregated loci across all datasets; mappable intra-chromosomal locus pairs are detected an average of 7-10 times within 10 Mb (**Extended Data Fig. 1d**), and 98.8-99.9% are found at least once at all genomic distances (**Table 1**).

**Table 1.** Summary of GAM datasets used in this study. VTA DNs were selected based on immunofluorescence detection of tyrosine hydroxylase (TH) from two animals, an 8-week old wild-type mouse and a 10-week old mouse carrying a TH- GFP reporter. PGNs were selected based on their position within the CA1 pyramidal cell layer and detection using pan-histone as a nuclear marker, from two 8-week old wildtype littermate mice. Cortical OLGs were selected based on detection of GFP expression from a 3-week old Sox10-cre-LoxP-GFP mouse. GAM data from mESC (clone 46C) was previously published^9^, and available from the 4DNucleome portal after quality control (https://data.4dnucleome.org/; **Supplemental** Table 6).

| Dataset | GAM samples collected (3NPs per sample) | GAM samples passing Quality Control | Number of cells per dataset | Samples that did not pass quality control |  |  | Water controls | % loci pairs detected at least once (50-kb resolution) |
| --- | --- | --- | --- | --- | --- | --- | --- | --- |
|  |  |  |  | Orphan windows >70% | Uniquely mapped reads <50,000 | Cross-contaminated |  |  |
| **DNs R1** | 656 | **585** | **1755** | 58 | 11 | 4 | 13 | 99.7 |
| **DNs R2** | 316 | **291** | **873** | 19 | 6 | 3 | 6 | 99.5 |
| **PGNs R1** | 218 | **209** | **209** | 7 | 2 | 0 | 2 | 99.9 |
| **PGNs R2** | 288 | **275** | **275** | 7 | 1 | 0 | 6 | 99.7 |
| **OLGs** | 336 | **290** | **290** | 46 | 4 | 3 | 0 | 98.8 |
| **mESCs** | - | **249** | **747** | - | - | - | - | 99.9 |

### ImmunoGAM detects 3D chromatin contacts of specific brain cell types

To explore differences in chromatin contacts for each cell type and biological replicate, we produced normalized chromatin contact matrices which inform about inter-locus 3D physical distances^3^. Visual inspection of contact matrices shows clear differences in pairwise contacts between all four cell types, with PGNs and DNs having the highest similarity, which is recapitulated in the biological replicates (**Fig. 1c**). For example, cell-type specific 3D genome topologies are observed across a 5- Mb region centered on the *Htr1a* gene, which encodes the 5-hydroxytryptamine (serotonin) receptor 1A. HTR1A modulates synaptic transmission in the hippocampus, and has single-nucleotide polymorphisms (SNPs) associated with schizophrenia^15^. *Htr1a* is not expressed in mESCs nor in OLGs, and is embedded in strongly interacting domains that have different conformations in DNs and in PGNs (**Fig. 1c**). An additional example on chromosome 17 shows strong contact patches that span 35 Mb in the three brain cell types but not in the mESCs (**Fig. 1d**), and which are reproduced in the biological replicates (**Extended Data Fig. 1e**). Thus, immunoGAM reveals extensive short- and long-range rearrangements of the 3D chromatin architecture in neurons and OLGs in the adult mouse brain.

### Topological domains have cell-type specific boundaries that contain cell specialization genes

To investigate cell-type specific 3D genome topologies and how they relate with gene expression, we analyzed published single-cell transcriptomes from the cortex, hippocampus and midbrain after identifying the populations of cells equivalent to the brain cell types captured by immuno-GAM. We selected eight subtypes of mature OLGs and OLG progenitors^16^, three subtypes of PGNs from CA1^17^ and four DN subtypes from the VTA^18^ (**Extended Data Fig. 2a**). We also generated single cell transcriptome data from mESCs and showed that it correlates with published bulk RNA-seq data^19^(**Extended Data Fig. 2a-b**). Separation of the different transcriptomes into cell-type specific clusters was confirmed by Uniform Manifold Approximation and Projection (UMAP) dimensionality reduction, and validated by overlaying the expression of known cell-type marker genes (**Extended Data Fig. 2c-d**). Next, we produced pseudobulk expression datasets for each cell type by pooling single-cell transcriptomes. A threshold of regularized log (R-log) ≥2.5 was used to designate expressed genes (see **Methods**, **Extended Data Fig. 2e**). Examples of expression profiles for known marker genes for each cell type are shown in **Extended Data Fig. 2f.**

Next, we mapped the organization of topologically associating domains (TADs) in the four cell types, using the insulation score method^3, 20^. Extensive reorganization of TADs was found across the genome, for example, at the *Gabr* gene cluster, encoding GABA receptors which are not expressed in mESCs and are most active in PGNs, especially *Gabra2* and *Gabrb1* (**Fig. 2a**), and at the *Hist1* locus which contains clusters of replication-dependent histone-1 genes highly expressed in mESCs (**Extended Data Fig. 3a**). We find that TADs have similar average lengths (∼ 1Mb; **Extended Data Fig. 3b**), but their genomic positions vary extensively between cell types (**Fig. 2b** shows the top seven combinations; for all combinations see **Extended Data Fig. 3c**). One third of all TAD borders are unique to one cell type (7- 10% for each cell type), 8% are shared between the three brain cell types, but only 14% are common to all cell types (for example, only 29% of mESC TAD borders are found in the other brain cell types; **Fig. 2b**). The two neuronal cell types show only 35% conservation between TAD borders, in contrast with 59-65% conservation between their biological replicates (**Extended Data Fig. 3c-d**). We also find that TAD boundaries common to all cell types show stronger insulation in the brain cell types than mESCs, possibly reflecting structural domain stabilization upon terminal differentiation (**Fig. 2c**). In contrast, cell-type specific boundaries are less insulated, except for the borders that contain expressed genes in each brain cell type.

**Figure 2.**
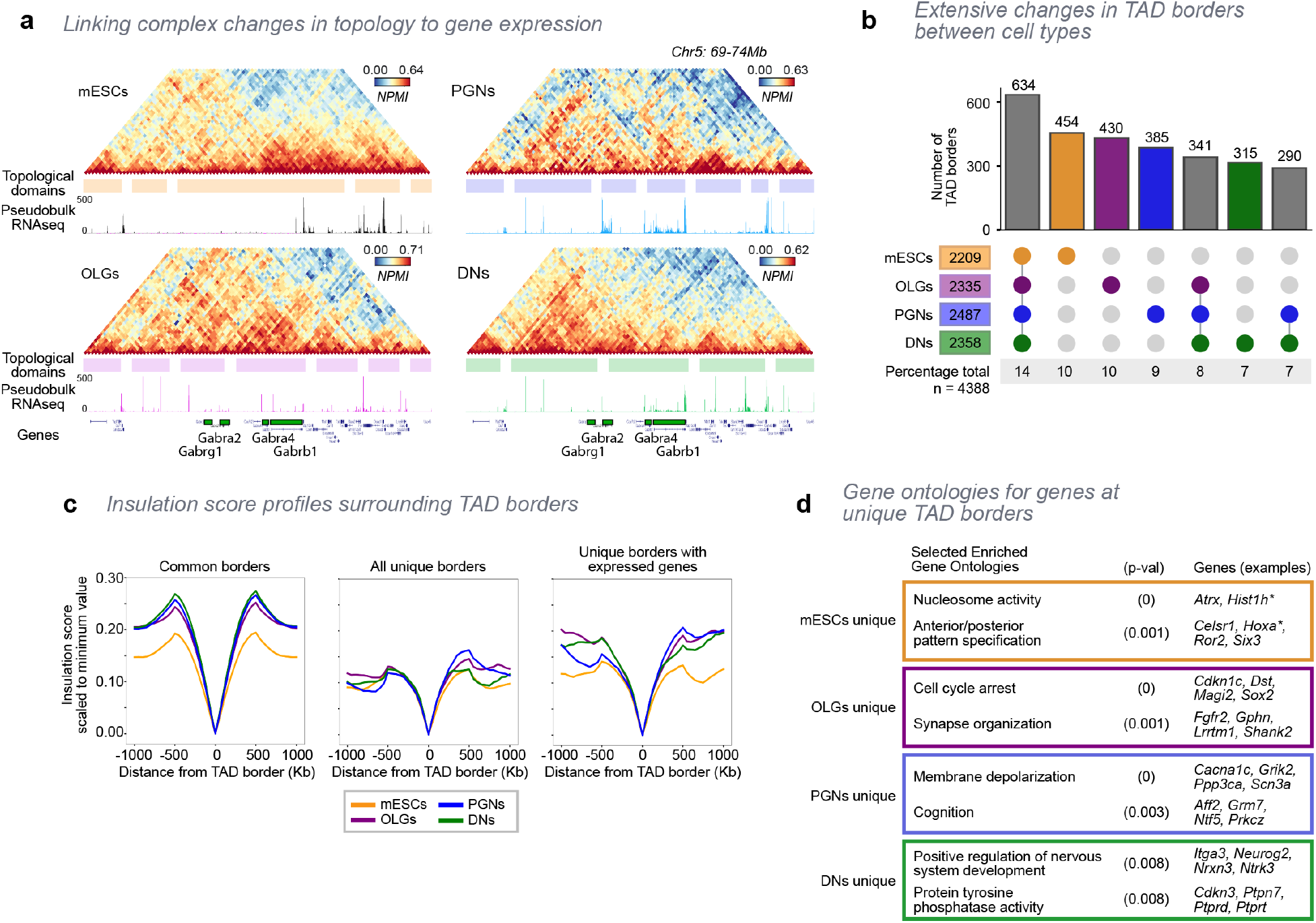
Extensive reorganization of topological domains in specialized cells of the brain. **a**, GAM contact matrices show cell-type specific differences in local contacts and topological domains within a 5-Mb region containing a cluster of GABA receptors (50-kb resolution). Shaded regions indicate topological domains. Mouse replicate 1 is shown for brain cells. **b**, Only 634 (14%) of borders are common to all cell types, followed by ∼315-454 borders that are cell-type specific. The top seven combinations are shown (for all combinations see **Extended Data Fig. 3c**). TAD boundaries were defined as 150-kb genomic regions centered on lowest insulation score window. **c**, Average insulation score profiles of common or unique borders centered on the lowest insulation point within each TAD border. Common TAD borders are more insulated in brain cell types than mESCs (*left*). Unique borders have similar boundary strength across cell types (*center*). Unique borders containing expressed genes are stronger in brain cell types than in mESCs (*right*). **d**, Selected gene ontologies for genes found at unique TAD borders. For all cell types, genes at unique TAD borders have enriched ontologies relevant for their cell functions (permuted p-values <0.05, 2000 permutations; Z-Score >1.96).

Next, we asked whether the rewired TAD boundaries contained cell type- specific genes. Genes found at unique TAD boundaries are enriched for gene ontology (GO) terms relevant for the specialized functions of each cell type (**Fig. 2d**). mESC-specific boundaries are enriched for genes associated with ‘nucleosome activity’ (e.g. *Hist1* locus), and ‘anterior/posterior pattern specification’ (e.g. *Hoxa* genes involved in embryonic development^21^). Boundaries unique to OLGs are enriched for genes involved in ‘cell cycle arrest’, which relates with the recent terminal differentiation of many OLGs, ‘synapse organization’, including genes that encode synaptic proteins and neurotransmitter receptors expressed in OLGs, and with functions in the response to neuronal input and modulation of myelin formation^22^. PGN-specific TAD borders also contain cell specialization-related genes, with roles in ‘membrane depolarization’ and ‘cognition’, including the modulation of the response to neuronal activation^23^. DN-specific borders contain genes related to neuronal development, and notably ‘protein tyrosine phosphatase activity’, important for dopaminergic differentiation, regulation of tyrosine hydroxylase activity and dopamine synthesis, and G-protein coupled receptor mediation of dopaminergic cell signalling^24, 25^. We overlapped expressed genes with cell-type specific TAD boundaries, and found that 38% of mESC-specific TAD boundaries contain mESC expressed genes, while 52-55% of boundaries specific to the brain cell types contain expressed genes respective to each cell type (**Extended Data Fig. 3e**). Our results show that the mapping of TADs in pure populations of terminally-differentiated cells detects a previously unappreciated variation in TAD organization, which is strongly connected with the presence of cell-type specific genes at TAD borders.

### Extensive reorganization and loss of contact density of the Neurexin-3 gene in DNs

Many neuronal genes which are involved in specialized cell processes, such as synaptic plasticity, often are longer than 250kb, contain several alternative promoters, and produce many isoforms due to alternative initiation sites and complex RNA processing^26^. The regulation of long genes is especially important for neuronal function, as their disruption or the presence of SNPs have been identified in neurodevelopmental or degenerative disorders^27, 28^. Transcription of many long neuronal genes is also sensitive to topoisomerase inhibition suggesting their expression is highly dependent on topological constraints^29^. Neurexin-3 (*Nrxn3*) is a 1.54Mb long gene highly affected by topoisomerase inhibition. *Nrxn3* encodes a membrane protein involved in synaptic connections and plasticity, and for which genetic variation is associated with alcohol dependence and autism spectrum disorder^27, 30^. In mESCs, *Nrxn3* spans two TADs and with an almost undetectable expression level (**Fig. 3a, Extended Data Fig. 4a**). *Nrxn3* is increasingly expressed in OLGs, PGNs and DNs. Remarkably, we found a general loss of TAD contact density throughout the *Nrxn3* gene in DNs, especially in the second TAD, a phenomenon that we report here as ‘TAD melting’.

**Figure 3.**
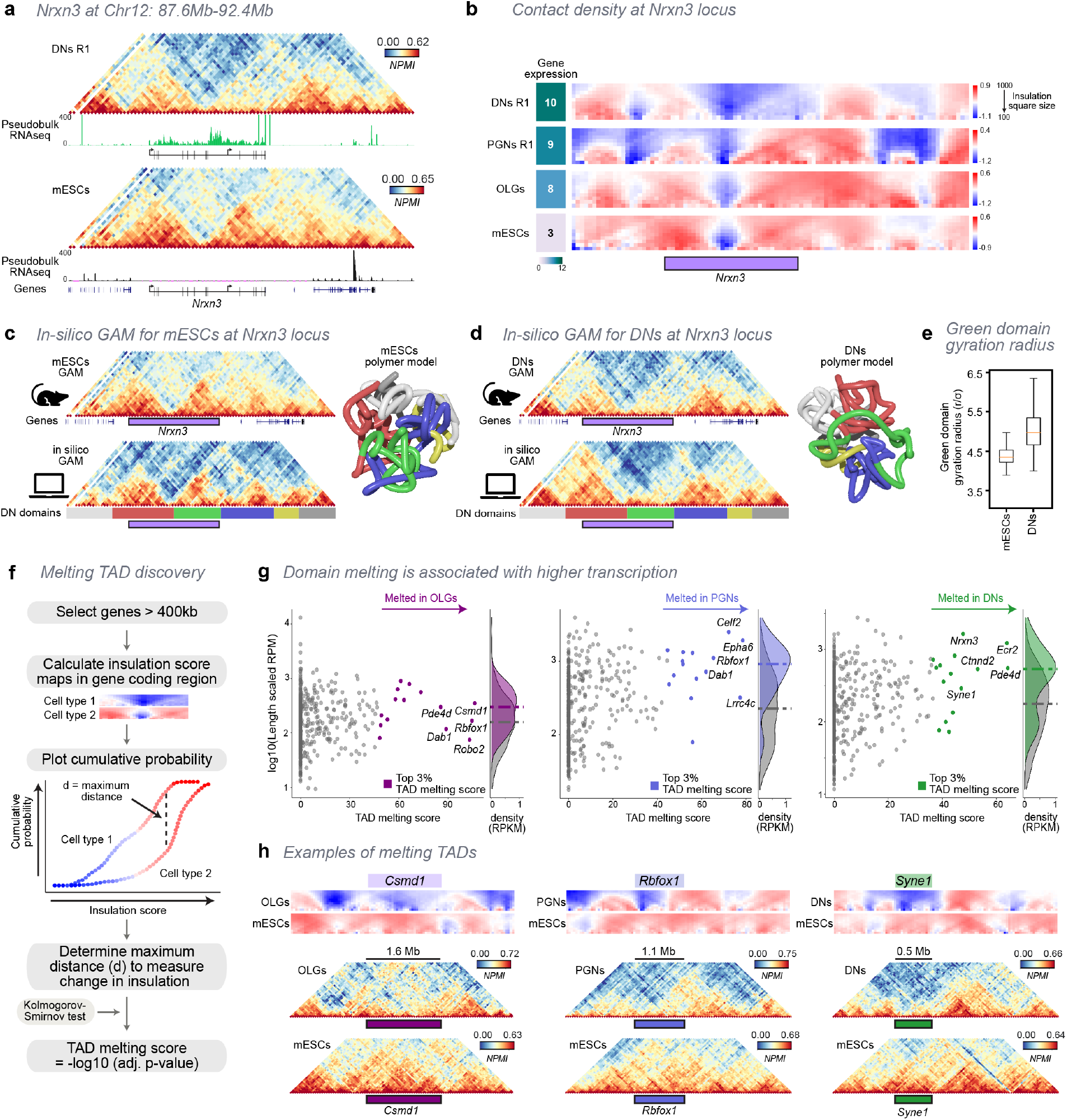
Extensive remodeling of chromatin structure and TAD melting at long and highly expressed genes. **a**, GAM contact matrices show the *Nrxn3* locus, a long (1.5Mb) gene which is highly expressed in DNs, but not in mESCs (50-kb resolution; chr12: 87,600,000-92,400,000). *Nrxn3* occupies two TADs with strong contacts in mESCs, which become decondensed, or ‘melted’ in DNs. **b**, Insulation score maps at the *Nrxn3* locus in mESCs, OLGs, PGNs and DNs (square sizes 100 - 1000 kb). *Nrxn3* contact density is strongest for mESCs, becoming increasingly decondensed in OLGs, PGNs and DNs, as its expression increases from R-log equal to 3, 8, 9 and 10, respectively. **c**,**d**, Ensembles of polymer models were produced for the *Nrxn3* locus in mESCs (**c**) and in DNs (**d**) from experimental GAM data using PRISMR modelling. The quality of the models was verified by applying in-silico GAM to the ensemble of polymers and comparison between NPMI-normalized contact matrices from in-silico and experimental immunoGAM (Pearson r = 0.72 and 0.79 for mESCs and DNs, respectively). Colour bars below *in-silico* matrices (Fig. 3 *legend, cont.)* highlight the position of domains in DNs and are used to colour the polymer examples shown. *Nrxn3* locus polymers have a globular structure with subdomains that interact with one another in mESCs (**c**). In DNs, the second TAD containing the Nrxn3 gene (green domain) becomes extended and loops away (**d**). **e**, The second *Nrxn3* TAD (green domain) has higher gyration radii in DNs than mESCs, consistent with its decondensed, melted state in DNs. **f**, TAD melting was determined for long expressed genes (>400 kb, 479 genes) from insulation score heatmaps. The distributions of insulation score values were compared between brain cell types and mESCs. The maximum distance between the distributions was determined using one-sided Kolmogorov-Smirnov testing and a TAD melting score was calculated based on the distribution dissimilarity. **g**, Genes with the top melting scores have higher RPM values, especially in PGNs and DNs. **h**, Examples of genes with top melting scores in each brain cell type.

To compare the changes in contact densities between cell types, we visualized the heatmap of a range of insulation scores (**Fig. 3b**). We found strong contact density present in both TADs in mESCs and OLGs although *Nrxn3* has higher expression in OLGs compared to mESCs (R-log = 8 compared with R-log = 3). With further increases of *Nrxn3* gene expression in PGNs (R-log = 9), the contact density became weaker in the first TAD, before extensive loss of contact density in DNs (R- log = 10). Biological replicates in PGNs and DNs confirm the increased levels of melting in DNs (**Extended Data Fig. 4b**).

### Modelling the loss of Nrxn3 TAD structure shows decompaction of chromatin

To better understand how the loss of TAD structure could influence 3D chromatin folding, we used a polymer-physics-based approach (PRISMR)^31^ to model the 3D structure of the *Nrxn3* region in mESCs and DNs (**Fig. 3c-d**). We generated 3D models from the GAM matrices, which we validated by applying an “*in-silico* GAM” approach^10^ to randomly section a single nuclear profile for each model. Reconstructed *in-silico* GAM chromatin contact matrices closely resemble the experimental data for both mESCs (Pearson r = 0.72) and DNs (r = 0.79). Importantly, the loss of contacts in the second *Nrnx3* TAD is captured by the *in-silico* matrices.

Next, we inspected single polymer models (**Fig. 3c,d**; coloured according to the structure of domains found in DNs). In the mESC models, the structure of the region was globular and highly intermingled (**Supplemental movie 1**; additional example models for both cell types can be found in **Extended Data Fig. 4c**). The second *Nrxn3* TAD (in green) interacts frequently with the surrounding domains. Remarkably, the second *Nrxn3* TAD is highly extended in DNs (**Supplemental movie 2**). Across the ensemble of DN polymer models, the second *Nrxn3* TAD shows much higher gyration radii compared to the same genomic region in mESCs, indicating that the region tends to be highly decondensed or ‘melted’ (**Fig. 3e**). In contrast, the TADs upstream and downstream to *Nrxn3* show marked decreases in gyration radii in DNs, which may be related to changes in gene expression in the neighbouring domains and/or potentially to torsional stress due to extensive *Nrxn3* transcriptional activity (**Extended Data Fig. 4d**). The loss of contact density and decondensation observed in *Nrxn3* highlights the complicated restructuring of local and long-range chromatin organization for different activation states of long neuronal genes, which can result in loss of the topological structure, decompaction, and melting of entire TADs.

**Figure 4.**
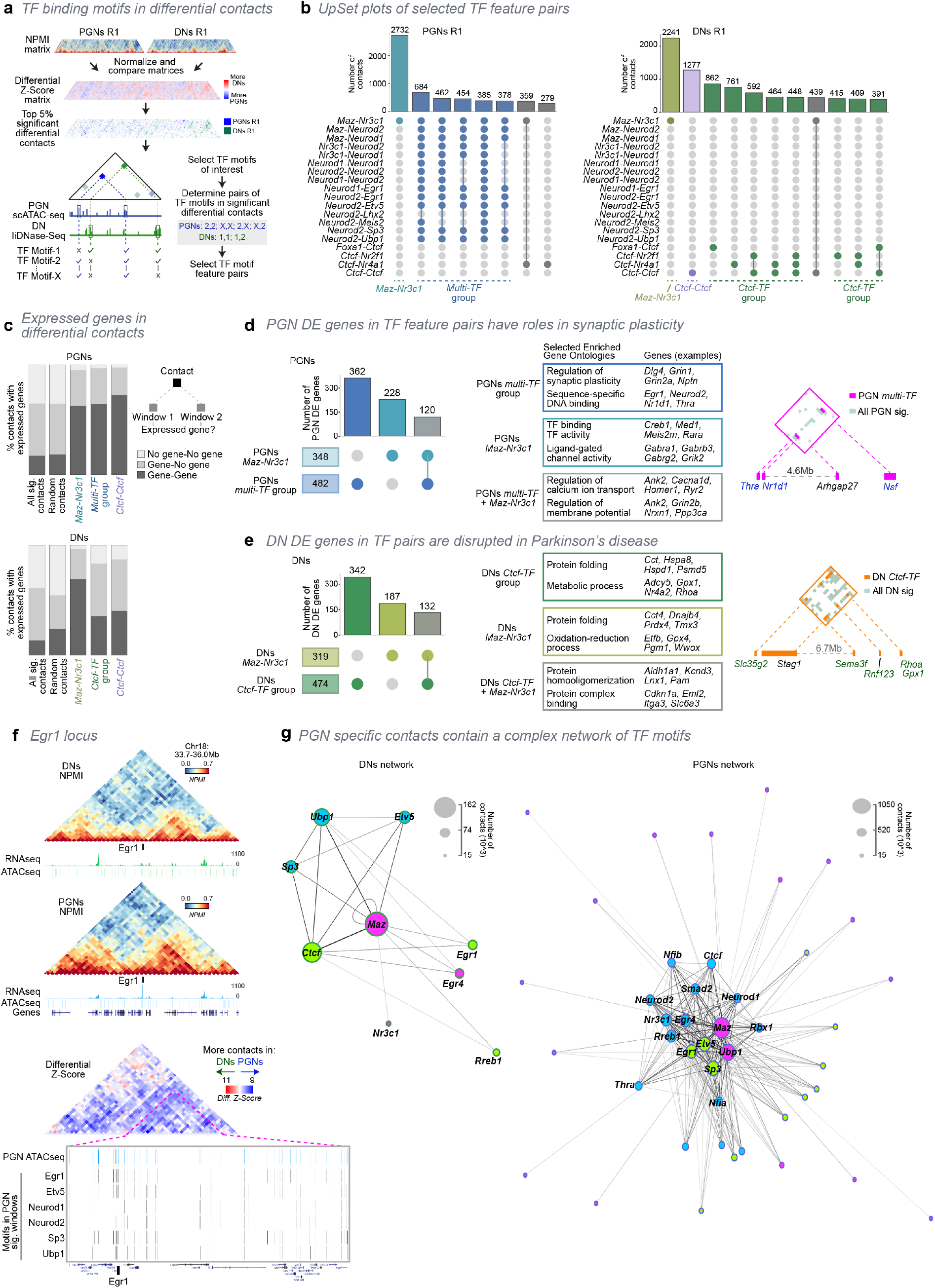
Neuron-specific genes establish PGN- and DN-specific contacts, which contain rich networks of TF binding sites within accessible regions. **a**, GAM contacts from PGNs and DNs were normalized and subtracted to produce a matrix of cell-type differential contacts (differential Z-Score matrix). The top 5% differential contacts for PGNs or DNs within 10 Mb were selected, and overlapped with accessible chromatin (Fig. 4 *legend, cont.)* regions to identify candidate regulatory elements. These accessible elements were used to search for putative binding sites of PGN and DN differentially expressed transcription factors (TFs). The most abundant TF feature pairs (top 20) were further investigated. R1, mouse replicate 1. **b**, Multiple TF feature pairs coincide in the same PGN (*left*) or DN (*right)* differential contacts. The top groups of contacts are shown for each cell type. **c**, Differential contacts with the most enriched TF feature pairs in PGNs and DNs contain expressed genes in at least one contacting window, often in both. **d**, Differential contacts involving the most abundant TF feature pairs in PGNs contain differentially expressed genes with PGN-specific roles. (*Right*) Matrix shows differential contacts established between PGN upregulated genes. Gene names highlighted in blue are upregulated in PGNs. **e**, Differential contacts containing the most abundant TF feature pairs in DNs also contain differentially expressed genes with roles in DN-specific functions. (*Right*) Matrix shows contacts between DN upregulated genes. Gene names highlighted in green are upregulated in DNs. **f**, In DNs, the *Egr1* gene establishes weak contacts with a large downstream domain. In PGNs, *Egr1* is upregulated and shows stronger interactions within the downstream domain. The differential contact (Z-Score) matrix shows increased PGN-specific contacts in the whole region surrounding *Egr1*. The *Egr1*-containing TAD has multiple TF binding sites found within PGN accessible regions. **g**, Network analysis and community detection for TF motifs found within DN or PGN differential contacts.

### Genome-wide detection of TAD melting is related with high transcription of long genes

To investigate whether TAD melting occurred in other long genes and in all brain cell types, we devised a pipeline to detect TAD melting based on the differences between cumulative probabilities of insulation scores. We determined a ‘TAD melting score’, as the significance value of the differences between the maximum distances of cumulative probabilities between two cell types **(Fig. 3f**). We calculated TAD melting scores for all genes longer than 400kb and expressed in at least one cell type (479 genes; **Supplemental Table 3**). We found that *Nrxn3* has TAD melting scores of 48 and 49 (replicates 1 and 2, respectively) when comparing DNs to mESCs (**Extended Data Fig. 4e**). We calculated TAD melting scores for all the brain cell types in comparison to mESCs, and explored further the genes with the top 3% of TAD melting scores in each cell type (15 genes in each cell type; **Fig. 3g; Extended Data Fig. 4f** shows the results for PGN and DN biological replicates).

To investigate whether TAD melting is generally related with transcriptional activity, we compared melting scores to the length-normalized number of reads covering both exons and introns (length-scaled Reads per Million; RPM; see **Methods**). Calculating transcriptional activity over the whole gene body allowed us to capture intronic reads which were abundantly detected in many long genes, for example at the *Nrxn3* gene in DNs (**Fig. 3a**). We find that the highest melting scores are associated with highest transcription in all three brain cell types, most pronounced in PGNs and DNs (**Fig. 3g**). Remarkably, 55% of all genes with top melting scores in any cell type or replicate (24 of 44 genes) were previously found to be sensitive to topoisomerase inhibition^29^, compared to only 16% of the other genes tested (69 of 435; Fisher’s exact p-value = 4x10^-8^; **Extended Data Fig. 4g**).

Examples of several genes with the highest melting scores are shown in **Fig. 3h** (see **Extended Data Fig. 4h** for cumulative probability scores). In OLGs, the top melting score was found for *Csmd1*, which is sensitive to topoisomerase inhibition^29^ and a multiple sclerosis^32^ and schizophrenia risk gene^33^. In PGNs, *Rbfox1* is a top melting gene, also sensitive to topoisomerase inhibition, with mutations linked to epilepsy and autism spectrum disorder^34, 35^. In DNs, another example includes *Syne1*, with mutations leading to muscular dystrophy and Parkinsonian-like symptoms^36^. Together, these observations highlight that TAD melting is an appreciated topological feature robustly captured by immunoGAM, which occurs at very long genes, often when these genes have the highest expression levels.

### Cell-type specific chromatin contacts contain binding motifs of differentially expressed transcription factors

To explore whether the extensive rearrangements in chromatin topology found in specialized brain cells are related with changes in the landscape of cis-regulatory elements, we investigated in detail the differences in specific chromatin contacts for PGNs and DNs, and whether they contain putative binding sites for neuronal specific transcription factors (TFs). First, we determined differential chromatin contacts by extracting the top 5% cell-type specific contacts below 10Mb (**Fig. 4a**). We identified locations of TF binding motifs for 218 TFs expressed in PGNs and/or DNs. To restrict the search space within the 50-kb windows, we collected open chromatin regions from published scATAC-seq experiments in PGNs^37^ and from single-cell low-input chromatin accessibility (liDNAse-seq) peaks in DNs^38^. Next, we selected TFs that are differentially expressed in PGNs and DNs and with motifs present in at least 5% of differential windows, finding 16 DN-specific TFs and 32 PGN-specific TFs (**Extended Data Fig. 5a-d**). We searched for pairs of windows with TF motifs (across the 1176 possible pairwise TF combinations) that were most enriched in one or the other cell type, or with a high ability to distinguish contacts between each cell type (see **Extended Data Fig. 5e** for full pipeline and criteria). Twenty TF motif pairs were considered for further analysis, containing combinations of 14 of the initial 48 TF motifs considered (**Extended Data Fig. 6a**, see **Supplemental Table 4** for all TF pairs).

We next searched for the overlap between TF motifs in the same pairs of contacting windows (**Fig. 4b**). The most frequent pair of TF motifs in both cell types is *Maz-Nr3c1* (2732 and 2241 contacts in PGNs and DNs, respectively) which involves *Maz* (which targets NMDA receptor genes^39^) in one contact anchor, and *Nr3c1* (associated with depression and anxiety disorders^40^) in the other anchor (**Fig. 4b, Extended Data Fig. 6b**). In PGNs, the next five groups of contacts have multiple TF feature pairs that include *Maz*-*Nr3c1*, plus combinations of eight different TFs: *Neurod1, Neurod2, Egr1, Etv5, Lhx2, Meis2, Sp3,* and *Ubp1* (*multi-TF* group; combined total of 2363 contacts). In contrast, in DNs, the most abundant group of contacts after *Maz-Nr3c1*, are contacts that do not contain *Maz-Nr3c1*, and have instead *Ctcf-Ctcf* alone (*Ctcf-Ctcf* group; 1277 contacts) or multiple groups of *Ctcf* heterotypic contacts with *Foxa1*, *Nr2f1* or *Nr4a1* (*Ctcf-TF* group; combined total of 4343 contacts). The significant contacts in all groups are found across all genomic distances considered (50kb-10Mb) though slightly enriched at >3Mb (**Extended Data Fig. 6c**). In light of the association between CTCF and TAD borders^41^, we checked whether contacts overlap with TAD borders. We find that only ∼25-35% of contacts involve a TAD border, except for *Ctcf-Ctcf* contacts in DNs, of which 47% involve a TAD border (11% involve a TAD border at both windows; range of contact distances is 50kb-2Mb; **Extended Data Fig. 6d**).

### Cell-type specific chromatin contacts contain differentially expressed genes

Next, we investigated whether the genomic regions involved in cell-type specific contacts connect neuronal genes with their candidate regulatory regions or with other genes, and whether they share neuronal functions. Remarkably, we found that a high percentage of differential contacts within the TF motif groups contain expressed genes in both contacting windows (36-72%, 25-47% in only one window; **Fig. 4c**). Many of these genes are differentially expressed (1609 and 2149 genes, in PGNs and DNs respectively), except for the *Ctcf-Ctcf* group which has the lowest number of differentially expressed genes (**Extended Data Fig. 6e**).

Many of the differentially expressed TFs with highest coverage in the PGN and DN differential contacts are known to have roles in neuronal activation^42, 43^, therefore we focused our further analyses on the genes most highly expressed in the cell type where they have stronger contacts. In PGN-specific contacts, the PGN-upregulated genes are found in the *multi-TF* (362 genes) or *Maz-Nr3c1* (228) contacts, with 120 genes sharing both types of contact (**Fig. 4d**). These genes share GO terms related to synaptic plasticity, the regulation of calcium transport, and TF binding and activity.

Interestingly, several of the genes are themselves TFs with motifs enriched in the PGN-specific contacts, including *Egr1* and *Neurod2*, also involved in the response to neuronal activation and synaptic plasticity^42, 43^. Two other genes encoding for PGN-upregulated TFs, *Thra* and *Nr1d1*, are closely located in chromosome 11 (separated by ∼3 kb) and establish a hub of *multi-TF* contacts with *Arhgap27* (at 4.6 Mb distance), involved in cognitive function^44^, and *Nsf* (5.1 Mb), which binds AMPA receptors and regulates both long-term potentiation (LTP) and depression (LTD; **Extended Data Fig. 6f** shows Z-Score matrix)^45^.

DN-upregulated genes are exclusive to either the *Ctcf-TF* (342) or *Maz-Nr3c1* groups, with 132 genes sharing contacts with both groups (**Fig. 4e**). Their GO terms are related to protein folding and metabolic functions, with many proteins in these groups dysregulated in Parkinson’s disease (PD) and other neuronal disorders. For example, *Gpx1* and *Rhoa* are both involved in oxidation-reduction reactions and disrupted in PD^46, 47^, and make *Ctcf-TF* containing contacts with the DN-upregulated genes *Semf3* (involved in neural circuit formation and a PD risk gene^48^), and *Rnf123* (an E3-ubiquitin ligase implicated in major depressive disorder^49^). These four genes form a hub of specific *Ctcf-TF* containing contacts with the schizophrenia risk gene *Slc35g2*^50^ and the cohesin subunit *Stag1* (**Extended Data Fig. 6g** shows Z-Score matrix).

We explored further *Egr1*, an immediate early gene upregulated in activated neurons^42^, which establishes PGN-differential contacts that contain *Egr1* and *multi-TF* motifs. In DNs, the *Egr1* gene is expressed (R-log = 6.8), and found at a TAD border with some interactions with its right-sided TAD (**Fig. 4f, Extended Data Fig. 6h**). In PGNs, where cells are frequently activated^12, 13^, *Egr1* is highly upregulated (R- log = 9.5) and becomes strongly embedded with the right TAD. In the differential contact matrix, most contacts within the *Egr1*-containing TAD are stronger in PGNs and associated with a high density of ATAC-seq peaks that contain complex combinations of motifs for the TFs found in the group of contacts containing the *Egr1* gene (sixth group in PGNs; **Fig. 4b**), including *Egr1, Etv5*, *Neurod1, Neurod2, Sp3* and *Ubp1*.

### PGN-specific contacts contain complex interconnected networks of TF binding sites

To explore the complex overlap of multiple TF motifs observed in the PGN- but not the DN-differential contacts more broadly, we next explored the degree of interconnectivity of TF motifs in differential contacts without pre-selection of the windows containing specific TFs (i.e. across the 1176 possible pairwise combinations of the 48 TFs). We built networks of all 48 TF motifs based on their co-occurrence in the top 5% differential contacts of PGNs or DNs, followed by community detection using the Leiden algorithm (see **Methods**, **Extended Data Fig. 6i, Supplemental Table 5**). Each TF motif was a node of the network, and the number of differential contacts between windows containing the motifs were the edges. To detect the strongest patterns, we considered only the TF motifs involved in >15,000. In DNs, we detected three communities with *Maz* as the central motif for one community, which also contained *Nr3c1* and *Egr4* (**Fig. 4g**). *Ctcf* is the central motif in a second community which also containing *Egr1* and *Rreb1*, and *Ubp1* is central for a third community, with *Sp3 and Etv5*. The three communities are inter-connected, with the total number of contacts for each motif ranging from ∼15,000-162,000. In striking contrast, the PGN network has a dense core of highly connected TF motifs, with up to ∼1,040,000 total contacts for *Maz*, which is also the central motif for one community. Two other communities are detected, with *Egr1* and *Etv5* as central motifs for one community, and several core TF motifs, including *Egr4, Neurod1, Neurod2, Rerb1* and *Smad2*, as central in the other community. All three communities were interconnected at the core of the network, while more peripheral TFs interact strongly with the core of its community. Together, these results suggest that chromatin contacts specific for different neuron types contain inter-connected hubs of TF motifs, which describe differentially expressed genes involved in specialized cell functions.

### Extensive A/B compartment changes occur between brain cells and mESCs, and relate with changes in gene expression

As a final exploration of chromatin contacts in mESCs and the three brain cell types, we investigated large-scale changes of compartment domains associated with open and closed chromatin by performing principal component analyses (PCA)^3^.

Visual comparisons of normalized compartment eigenvectors show extensive variation of compartments A and B between mESCs and the three brain cell types (**Fig. 5a**, **Extended Data Fig. 7a**). The compartment eigenvector values are best correlated between biological replicates, next between DNs and PGNs, neurons and OLGs, and least between the different brain cells and mESCs (**Extended Data Fig. 7b**). A large fraction of windows were in compartment A (40%, 2325 windows) in all cell types, but only 15% of windows share compartment B membership (898 windows; **Fig 5b, Extended Data Fig. 7c-d**). The next most abundant group of windows undergo changes between compartments B in mESC to A in all the brain cell types (12%, transition B-mESC → A-brain cells) or between compartment A in mESCs to B in brain cells (7%, B-mESC → A-brain cells). Regardless of the changes in compartment membership, the mean and total genomic lengths occupied by A or B compartments is similar across cell types (**Extended Data Fig. 7e-f**).

**Figure 5.**
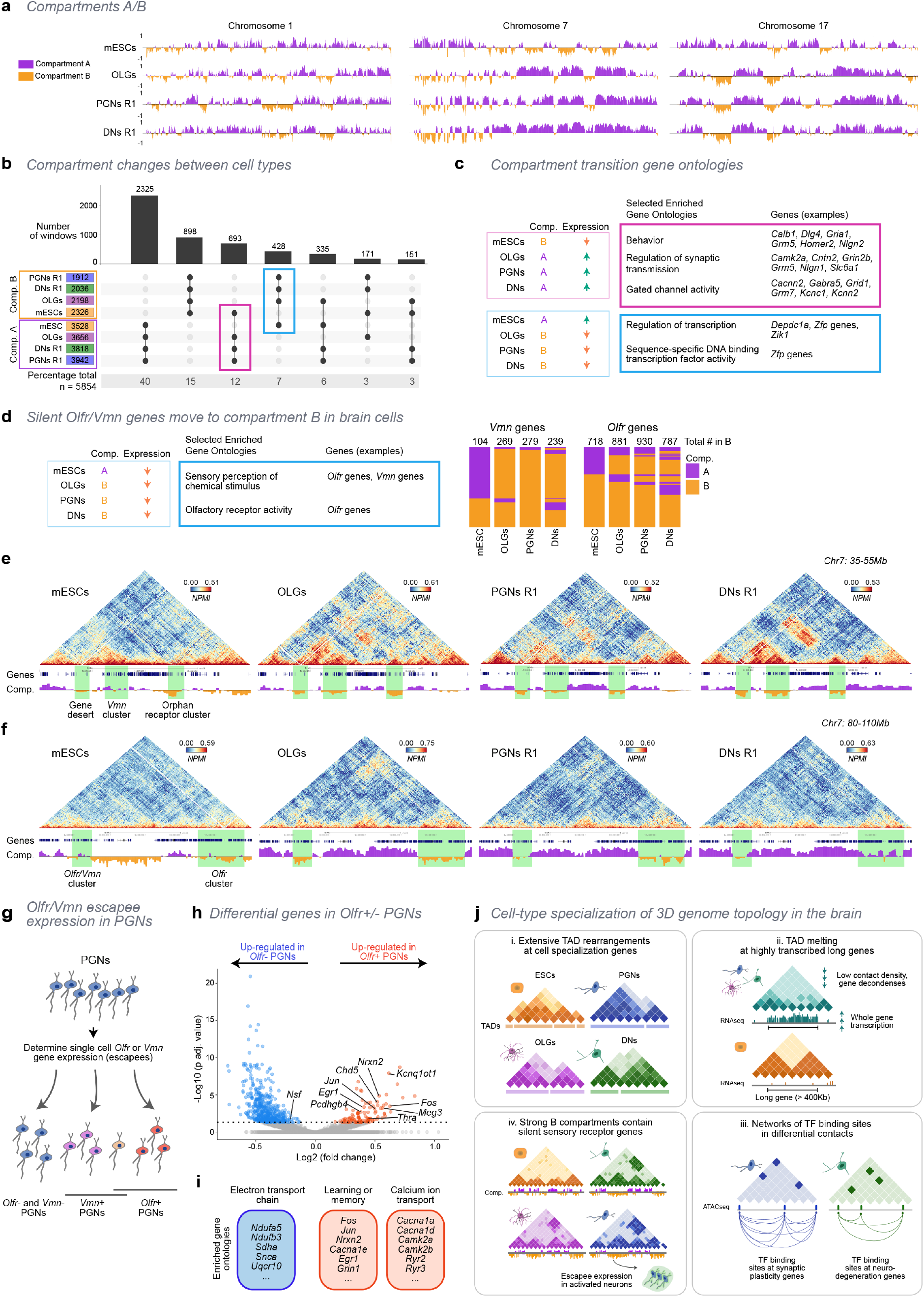
Clusters of sensory receptor genes more often belong to B compartments in brain cells and establish megabase-range interactions. **a**, Open and closed chromatin compartments (A and B, respectively) display different genomic distributions in mESCs, OLGs, PGNs and DNs. Normalized PCA eigenvectors were (Fig. 5 *legend, cont.)* computed from GAM datasets from mESCs and for brain cell types, and used to classify compartments. Mouse replicate 1 (R1) is shown for PGNs and DNs. **b**, Most genomic windows share membership to compartments A, followed by B in all cell types. The most frequent compartment changes occur from compartment B in mESCs to A in all brain cells (pink box), followed by from A in mESCs to B in all brain cells (blue box; for all combinations see **Extended Data** Fig. 7d). **c**, Selected gene ontologies (GOs) for genes that increase expression in all brain cells relative to mESCs and move from compartment B in mESCs to compartment A in brain cells (*pink box*); and for genes that decrease expression in brain cells and move to compartment B, compared to mESCs (*blue box*). All enriched GOs had permuted p-values = 0. **d**, Selected GOs for genes that remain silent in all cell types but gain membership to compartment B in brain cells. Most genes are *Olfr* and *Vmn* sensory receptor cluster genes. A higher proportion of *Olfr* and *Vmn* genes are found in B compartments in brain cells, compared to mESCs. All enriched GOs had permuted p-values = 0. **e**, GAM contact matrices containing *Vmn* and orphan receptor genes shows strong interactions separated by 10 Mb between B compartments in OLGs, PGNs and DNs, but not mESCs. Shaded green regions indicate interacting regions. **f**, GAM contact matrices show interactions between an *Olfr/Vmn* gene cluster and a second *Olfr* cluster (green shaded regions) separated by 25 Mb. The contacts between the two receptor clusters are strongest in OLGs, where the B compartment is strongest. **g**, *Olfr* or *Vmn* escapee expression in single PGN cells. PGN cells were classified as *Olfr+* or *Vmn+* if at least one *Olfr* or *Vmn* gene was detected, respectively. **h**, Volcano plot of differentially expressed (DE) genes between Olfr-expressing and non-expressing PGNs. Genes with an adjusted p-value < 0.05 were considered DE. **i**, Selected gene ontologies (GOs) of upregulated genes were enriched for neuronal activation functions, while downregulated genes are involved in mitochondria metabolism. **j**, Summary diagram. The 3D genome is extensively reorganized in brain cells. (i) TAD borders are rearranged, with new border formation coinciding with genes important for cell specialization in all cell types. (ii) TAD melting occurs at very long and highly transcribed genes in brain cells. (iii) The most specific contacts in neurons contain complex networks of binding sites of neuron-specific transcription factors. Contacts bridge genes upregulated in the neurons where the contacts are observed. Genes in contact hubs have functions in synaptic plasticity (PGNs) and neurodegenerative disease (DNs). (iv) Finally, B compartments contain clusters of sensory receptor genes silent in all cell types which form strong megabase-range contacts. A subset of sensory receptor genes escape repression when neurons are activated.

To investigate how compartment changes between mESCs and the brain cell types related to gene expression, we clustered genes found in each type of transition according to their expression in each cell type. We find 335 genes within the regions with transition B-mESC → A-brain cells, which are more strongly expressed in all the brain cell types than in mESCs (**Extended Data Fig. 8a**). These genes share functions between the brain cell types and have enriched GO terms such as “behavior”, “regulation of synaptic plasticity” and “gated ion channel activity” (**Fig. 5c**). In regions that undergo A-mESC → B-brain cells transitions, we remarkably find that most genes are silent in all cell types (572/715 genes that undergo A-to-B transitions), and only 50 genes are more expressed in mESCs (**Extended Data Fig. 8b**). Enriched GO terms for these 50 genes include transcriptional regulation and transcription factor activity, and include many *Zfp* genes, encoding zinc-finger binding proteins with roles in the epigenetic regulation of pluripotency factors, and silencing of active genes during the exit from pluripotency^51^.

### Clusters of sensory receptor genes are more strongly associated with B compartments in brain cells and establish megabase-range interactions

We were intrigued by the large number of genes that undergo A-mESC to B-brain-cell transitions irrespectively of being silent in all cell types. This group of genes contained sensory receptor genes such as *Vmn* (vomeronasal; 149 genes out of 572 silent genes in the group) and *Olfr* (olfactory receptor; 179 genes), which are most often found in gene clusters^52, 53^ (**Fig. 5d**). Although silent in mESCs, only 35% of *Vmn* genes (104/290) and 66% of *Olfr* genes (718/1095) are associated with compartment B, compared to 82-96% and 72-85% in brain cells, respectively (**Fig. 5d**).

We explored how the stricter compartment B association of sensory receptors in brain cells was related with chromatin contact formation in regions containing *Vmn* and *Olfr* clusters. Visual inspection of matrices and compartments on chromosome 7 (which contains the most *Vmn* and *Olfr* genes combined) showed that *Vmn* genes and/or *Olfr* genes are often involved in strong long-range patches of contacts only observed in brain cells, which can be separated by distances above 10 Mb and up to 50 Mb (**Fig. 5e,f**, **Extended Data Fig. 9a**). For *Olfr* clusters in **Fig. 5f**, the contacts between the two receptor clusters are most clearly found in OLGs, which are strongly within the B compartment. Contacts are also found in PGNs, but not in DNs where the second cluster is in the A compartment. The long-range patches of contacts on chromosome 17 shown in **Fig. 1d** also contain both *Vmn* and *Olfr* genes.

To investigate the strength of long-range contacts between *Vmn* or *Olfr* genes in non-adjacent compartments, we normalized matrices using Z-Scores at each genomic distance to account for their occurrence at different genomic lengths, and compared the top 20% of Z-Scores at distances >3 Mb. The brain cell types show higher Z-Scores for long-range contacts between B compartment regions containing *Vmn* genes, and to a lesser extent *Olfr* genes, when compared to mESCs or the genome-wide B compartment distribution, with no difference for genes found in compartment A for any cell type (**Extended Data Fig**. **9b**). These results suggest that sensory genes are not only more locally compartmentalized in heterochromatic B compartments, but also form stronger contacts in 3D with other B compartments in brain cells, which suggests a stricter repression mechanism in brain cells.

### PGN subpopulations expressing Olfr genes have gene expression signatures of early neuronal activation

Tighter B compartmentalization of *Vmn* and *Olfr* genes may be important for their repression in the brain cell types. *Olfr* genes have been reported to be stochastically activated, and mis-expressed in several neurodegenerative diseases in cortical PGNs^54^. Additionally, *Olfr* expression significantly increases after pharmacological activation of glutamatergic neuronal firing^7^ (**Extended Data Fig. 9c**), suggesting that some *Olfr* genes can escape repression, dependent on cell state. Taking advantage of the large number of single cell transcriptomes available for PGNs^17^, we tested whether single PGNs had any *Olfr* or *Vmn* escapee gene expression (**Fig. 5g,** see **Methods**). We find that 23% (203/876 cells) and 19% (164) of single PGNs have at least one transcribed *Vmn* or *Olfr* gene, respectively.

To determine whether *Olfr* or *Vmn* escapee expression in PGNs was stochastic or related with functional differences between cell states within the PGN population, we measured differential gene expression between cells expressing at least one *Olfr* (*Olfr+* PGNs) or *Vmn* (*Vmn+* PGNs) gene and cells that expressed neither (*Olfr-* or *Vmn-* PGNs). Remarkably, we found 87 genes significantly upregulated in the PGNs expressing at least one *Olfr* gene, and 460 genes which were significantly downregulated (**Fig. 5h, Extended Data Fig**. **9d-e**). In contrast, we found no differentially expressed genes for random subsets of PGNs, and very few between the *Vmn+* and *Vmn-* PGNs, suggesting their escapee expression is stochastic (**Extended Data Fig. 9f**).

To further investigate whether the cells with escapee expression of *Olfr* genes might have a specific physiological state, we performed GO analyses of the upregulated genes in *Olfr*-expressing PGNs. Strikingly, and in accordance to the *Olfr* gene expression in neurons upon pharmacological activation^7^ (**Extended Data Fig. 9c**), we found genes related to neuronal activation (**Fig. 5i**). For example, genes involved in the early response to LTP are found in *Olfr+* PGNs, such as the upregulated genes *Egr1*, *Fos*, *Jun*, *Nrxn2*, and *Grin1*^55^, and the downregulated gene *Nsf*^45^. Significantly downregulated genes mostly included mitochondrial related processes, which are coupled to and finely tuned during neuronal activation^55^. Finally, we observed that *Olfr* escapee genes, but not *Vmn* genes, are more likely to escape repression if located in compartment A (**Extended Data Fig. 9g**), suggesting that the tighter B compartmentalization of *Olfr* genes might be important for their repression in brain cell types, especially in PGNs which are undergoing neuronal activation and likely chromatin structure reorganisation^7^.

## Discussion

Here, we adapted the GAM method to capture genome-wide chromatin conformation states of selected cell populations in the mouse brain that perform specialized functions. Using immunoGAM, we selected nuclear slices from specific brain cells by immunofluorescence prior to laser microdissection, and characterized the relationship between 3D chromatin organization and cell specialization in intact tissues. We discovered dramatic reorganization of chromatin topology which reflected cell function and specialization at multiple genomic scales, from specific contacts to compartment changes that spanned tens of megabases (**Fig. 5j**). For example, we found extensive restructuring of topological domains, with 76% of TAD borders being different in at least one cell type. Unique borders in each cell type contained genes relevant for cell specialization, which may be functionally necessary for their interaction with cis-regulatory elements and transcriptional activation^57^.

Of particular importance was the observation that the highest expression levels of very long genes in brain cells corresponds with a massive reorganization of local and long-range structure. We show that many highly transcribed long genes lose chromatin contact density, which coincides with the decondensation and unfolding of an entire domain, a phenomenon that we call ‘TAD melting’. Many long genes are regulated in a specialized manner in terminally differentiated cells, for example by the activity of topoisomerases^29^, by the presence of long stretches of broad H3K27ac and H3K4me1 which act as enhancer-like domains^58^, by the formation of large transcription loops^59^, or through repressive mechanisms such as DNA methylation^60^. The regulation of long genes is further complicated by intricate splicing dynamics^26^, which require highly adaptive responses based on neuronal activation state. In fact, for many of the genes we highlight, including *Nrxn3*, *Csmd1*, *Rbfox1* and *Syne1*, genetic variants are often associated with or directly result in neuronal diseases^27, 30, 33–36^. Thus, understanding how genome folding relates with the response to environmental challenges is increasingly important to further our understanding of the mechanisms of neurological disease.

Our results strongly support that prediction of functional or disease states in a cell type could be inferred from chromatin architecture. When we searched for TF binding motifs within accessible regions of specific chromatin contacts for a cell type, we found rich networks of putative binding sites for differentially expressed TFs. These networks connect hubs of differentially expressed genes with specialized functions in neurons. For example, DN-specific contacts contained heterotypic *Ctcf* binding motifs, present in contact hubs containing metabolic and protein folding genes. The accumulation of oxidative damage and disruption of ubiquitin-mediated protein folding are pathological hallmarks of Parkinson’s disease^46, 47, 61^. Therefore, future studies will be important to understand the relationship between DN-specific chromatin landscapes, disease-associated genetic variation and the regulation of these critical genes, with potential implications for the etiology of Parkinson’s disease.

An important contrast found in the PGN-specific contacts was the complexity of TF binding motif networks, which contained hubs differentially expressed genes with roles in synaptic plasticity. PGNs frequently respond to environmental cues during long-term potentiation (LTP) or depression (LTD), either by external stimuli or by stochastic and intrinsic activation^12, 13, 42, 45^. Strikingly, PGN-specific contacts containing TF genes involved in the early activation of LTP such as *Egr1* and *Neurod2*, which were also marked by accessible sites containing their own binding motifs, suggesting mechanisms of self-activation by which expression of a TF increases contacts of the TF with other regions containing its own binding sites. Activity-mediated waves of transcription accompany LTP, from immediate early genes, to secondary response genes, followed by long-term upregulation of synaptic proteins. Further, positive feedback loops of BDNF signaling at local synapses have been proposed as a mechanism of LTP induction^62^. Together with reports that *de novo* chromatin looping can accompany transcriptional activation^7^, our work suggests that coordinated transcription factor binding at apparent distant locations in the linear genomic scale, but in close contact due to the 3D chromatin landscape, may be critical for LTP induction.

The interplay of complex architecture and LTP related functions in PGNs was further highlighted by the observed activation of a subset of PGNs with *Olfr*-gene escapee expression. *Olfr* expression escape coincided with immediate early gene and synaptic plasticity-related gene expression. The otherwise tight repression of *Olfr* genes within strong B compartment regions, which form cis-contacts tens of mega- bases away, was especially noteworthy as *Olfr* genes can have non-sensory organ functions in neurons related to specificity of neuronal activation^7, 54^. *Olfr* genes are highly repressed in mature olfactory sensory neurons, where they form a large inter-chromosomal hub to regulate specificity of single *Olfr* gene activation^63, 64^. Our results expand these previous findings, highlighting that alternative long-range mechanisms of *Olfr* repression may also reflect their specialized roles in different cell types of the brain.

Finally, our work shows that immunoGAM is uniquely positioned to probe questions related to cell state and specialized function within the brain or in other complex tissues, as local tissue structure is maintained. ImmunoGAM also has the potential to be applied in multiple cell types within the same tissue while retaining the geographic positions of each cell, a prospect that can further deepen our understanding of coordinated interactions between cells in complex diseases. As immunoGAM requires only very small cell numbers to capture complex differences in genomic topology (∼1000 cells), it may also have prognostic value in highly precious clinical patient samples, and to provide critical insights into the etiology and progression of neurological disease. Collectively, our work revealed that cell specialization in the brain and chromatin structure are intimately linked at multiple genomic scales.

## Acknowledgments

The authors thank Sheila Q. Xie and Azhaar Ashraf for help processing midbrain samples, Thomas M. Sparks for developing the NPMI normalisation pipeline, Catherine Baugher for developing the TF motif analysis pipeline, and the Pombo lab members for helpful discussions. AP and AA acknowledge support from the Helmholtz Association (Germany). AP and MN acknowledge support from the National Institutes of Health Common Fund 4D Nucleome Program grant U54DK107977, and the Berlin Institute of Health (BIH). AP and LZR acknowledge support by the Deutsche Forschungsgemeinschaft (DFG; German Research Foundation) International Research Training Group (IRTG2403). AP acknowledges support from the Deutsche Forschungsgemeinschaft (DFG; German Research Foundation) under Germany’s Excellence Strategy – EXC-2049 – 390688087. GCB acknowledges European Union Horizon 2020/European Research Council Consolidator Grant (EPIScOPE no. 681893), Swedish Research Council (no. 2015- 03558; 2019-01360), Swedish Brain Foundation (no. FO2017-0075), Knut and Alice Wallenberg Foundation (grant 2019-0107), The Swedish Society for Medical Research (SSMF, grant JUB2019), Ming Wai Lau Centre for Reparative Medicine and Karolinska Institutet. GD and GA acknowledge support from the Austrian FWF through DK W1206 “Signal Processing in Neurons” and SFB F44 “Cell Signaling in Chronic CNS Disorders”, P25014-B24. MAU acknowledges funding by the Medical Research Council (UK) (U120085816) and a Royal Society University Research Fellowship. MN thanks support from CINECA ISCRA Grant HP10CYFPS5 and HP10CRTY8P, by computer resources at INFN and Scope at the University of Naples (MN). LW acknowledges the support of Ohio University’s GERB program. IH was supported by a Boehringer Ingelheim Fonds PhD fellowship, and ETT by an EMBO short-term fellowship (ASTF 336-2015). II-A was supported by a Long-Term Fellowship from the Federation of European Biochemical Societies (FEBS).

## Author Contributions

AP designed the concept for this work; WW, MM, AA, EJP and AP collected animal tissues; WW, AK, IH, MM, LS and RK produced GAM datasets; WW, AK, IH, MM and LS optimised the experimental protocol; WW, AK, II-A, CJT and EI developed computational pipelines for bioinformatics and QC analyses of GAM data; AK and IIA performed QC analyses of GAM data; WW and AK performed bioinformatics analyses of GAM data; DS and CJT developed the melting TAD analysis; DS performed the melting TAD analysis of GAM data; ETT performed mESC culture experiments; WWN and CJT developed the differential contact analysis; WWN performed differential contact analysis of GAM data; ETT and AAK produced single- cell RNAseq data; LZR and DS performed RNA-seq analysis; LZR performed differential gene expression analyses; SB optimized polymer modeling and performed PRISMR analysis; SB and AMC produced models for polymer modeling; AMC performed statistical analyses of polymer models; LF and FM performed in- silico GAM experiments; YZ performed TF motif finding enrichment, network, and community analyses; SAT supervised scRNA-seq experiments; MM, GA, GD, MU and GCB supervised animal tissue collection and provided animal samples; AP supervised GAM experiments and bioinformatics analyses; VF, AA, and AP supervised the scRNA-seq analysis; MN supervised the polymer modeling and in- silico GAM; LW supervised the TF motif and network analyses; WW, AK, IH, LZR, DS, MM, YZ, SB, AMC, MN, LW, GCB and AP contributed to the interpretation of the results; WW wrote the first draft of the manuscript; WW and IH designed the figures; WW, AK, IH, LZR, DS, YZ, SB, AMC, LW, and AP wrote the manuscript. All authors provided critical feedback and helped revise the manuscript. The authors consider WW, AK and IH to have contributed equally to this work. The authors also consider LZR and DS to have contributed equally to this work.

## Materials & Correspondence

Correspondence and material requests should be addressed to

## Methods

### Animal maintenance

Collection of GAM data from dopaminergic neurons was performed using one C57Bl/6NCrl (RRID: IMSR_CR:027; WT) mouse which were purchased from Charles River, and from one TH-GFP (B6.Cg-Tg(TH-GFP)21-31/C57B6) mouse, obtained as previously described^64, 65^. All procedures involving WT and TH-GFP animals were approved by the Imperial College London’s Animal Welfare and Ethical Review Body. Adult male mice of age 2–3 months were used. All mice had access to food and water *ad libitum* and were kept on a 12 h:12 h day/night cycle. C57Bl/6NCrl and TH-GFP mice received an intraperitoneal (IP) injection of saline 14 days or 24 h prior to the tissue collection, respectively, and they were part of a larger experiment for a different study.

Collection of GAM data from somatosensory oligodendrocyte cells was performed using Sox10::Cre-RCE::loxP-EGFP^66^ animals which were obtained by crossing Sox10::Cre animals^67^ on a C57BL/6j genetic background with RCE::loxP- EGFP animals^68^ on a C57BL/6xCD1 mixed genetic background, both available at The Jackson Laboratories. The Cre allele was maintained in hemizygosity while the reporter allele was maintained in hemi- or homozygosity. Experimental procedures for Sox10::Cre-RCE::loxP-EGFP animals were performed following the European directive 2010/63/EU, local Swedish directive L150/SJVFS/2019:9, Saknr L150 and Karolinska Institutet complementary guidelines for procurement and use of laboratory animals, Dnr 1937/03-640. The procedures described were approved by the local committee for ethical experiments on laboratory animals in Sweden (Stockholms Norra Djurförsöksetiska nämnd), lic.nr. 130/15. One male mouse was sacrificed at P21. Mice were housed to a maximum number of 5 per cage in individually ventilated cages with the following light/dark cycle: dawn 6:00-7:00, daylight 7:00-18:00, dusk 18:00-19:00, night 19:00-6:00.

Collection of GAM data from hippocampal CA1 pyramidal glutamatergic neurons was performed using two 19 week-old male *Satb2^flox/flox^* mice. C57Bl/6NCrl (RRID: IMSR_CR:027; WT) mice were purchased from Charles River, Satb2^flox/flox^ mice that carry the floxed exon 4 have been previously described^69^. The experimental procedures were done according to the Austrian Animal Experimentation Ethics Board (Bundesministerium für Wissenschaft und Verkehr, Kommission für Tierversuchsangelegenheiten). All mice had access to food and water ad libitum and were kept on a 12 h:12 h day/night cycle.

### Tissue fixation and preparation

WT, TH-GFP, and *Satb2*^flox/flox^ mice were anaesthetised under isoflurane (4 %), given a lethal IP injection of pentobarbital (0.08 μl; 100 mg/ml; Euthatal), and transcardially perfused with 50 ml of ice-cold phosphate buffered saline (PBS) followed by 50-100 ml of 4% depolymerised paraformaldehyde (PFA; Electron microscopy grade, methanol free) in 250 mM HEPES-NaOH (pH 7.4-7.6). Sox10::Cre-RCE::loxP-EGFP animals were sacrificed with a ketaminol/xylazine intraperitoneal injection followed by transcardial perfusion with 20 ml PBS and 20 ml 4% PFA in 250 mM HEPES (pH 7.4-7.6). From C57Bl/6NCrl or TH-GFP mice, brains were removed and the tissue containing the VTA were dissected from each hemisphere at room temperature, and quickly transferred to fixative. For *Satb2*^flox/flox^ mice, the CA1 field hippocampus was dissected from each hemisphere at room temperature. For Sox10Cre/RCE mice, brain tissue containing the somatosensory cortex was dissected at room temperature. Following dissection, tissue blocks were placed in 4% paraformaldehyde (PFA) in 250 mM HEPES-NaOH (pH 7.4-7.6) for post-fixation at 4°C for 1 h. Brains were then placed in 8% PFA in 250mM HEPES and incubated at 4°C for 2-3 h. Tissue blocks were then placed in 1% PFA in 250 mM HEPES and kept at 4°C until tissue was prepared for cryopreservation (up to 5 days, with daily solution changes).

### Cryoblock preparation and cryosectioning

Fixed tissue samples from different brain regions were further dissected to produce ∼1.5x3 mm tissue samples suitable for Tokuyasu cryosectioning^3^ (**Extended Data Fig. 1a**), at room temperature in 1% PFA in 250 mM HEPES. For the hippocampus, the dorsal CA1 region was further isolated. Approximately 1-3 mm x 1- 3 mm blocks were dissected from all brain regions and were further incubated in 4% PFA in 250 mM HEPES at 4°C for 1 h. The fixed tissue was transferred to 2.1 M sucrose in PBS and embedded 16-24 h, at 4°C, positioned at the top of copper stub holders suitable for ultracryomicrotomy and frozen in liquid nitrogen. Cryopreserved tissue samples are kept indefinitely immersed under liquid nitrogen.

Frozen tissue blocks were cryosectioned with a Ultracryomicrotome (Leica Biosystems, EM UC7), with an approximate thickness of 220-230nm (ref. ^3^). Cryosections were captured in drops of 2.1 M sucrose in PBS solution suspended in a copper wire loop and transferred to 10 mm glass coverslips for confocal imaging, or onto a 4.0 µm polyethylene naphthalate (PEN; Leica Microsystems, 11600289) membrane on metal framed slides for laser microdissection.

### Immunofluorescence detection for confocal microscopy

For confocal imaging, cryosections were incubated in sheep anti-tyrosine hydroxylase (TH, 1:500; Pel Freez Arkansas LLC, Cat#P60101-0), mouse anti-pan-histone H11-4 (1:500; Merck, Cat#MAB3422) or chicken anti-GFP (1:500; Abcam Cat#ab13970) followed by donkey anti-sheep or goat anti-chicken IgG conjugated with AlexaFluor-488 (for TH and GFP; Invitrogen) or donkey anti-mouse IgG conjugated with AlexaFluor-555 or AlexaFluor-488 (for pan-histone; Invitrogen).

For PGNs, cryosections were washed (3x, 30 min total) in PBS, permeabilized (5 min) in 0.3% Triton X-100 in PBS (v/v) and incubated (2 h, room temperature) in blocking solution (1% BSA (w/v), 5% FBS (w/v) (GibcoTM Cat#10270), 0.05% Triton X-100 (v/v) in PBS). After incubation (overnight, 4°C) with primary antibody in blocking solution, the cryosections were washed (3-5x; 30 min) in 0.025% Triton X- 100 in PBS (v/v) and immunolabeled (1 h, room temperature) with secondary antibodies in blocking solution, followed by 3 (15 min) washes in PBS. Cryosections were then counterstained (5 min) with 0.5 µg/ml 4’,6’-diamino-2-phenylindole (DAPI; Sigma-Aldrich^®^ Cat#D9542) in PBS, and then rinsed in PBS and water. Coverslips were mounted in Mowiol® 4-88 solution in 5% glycerol, 0.1M Tris-HCl (pH 8.5).

The number of SATB2 positive cells present in the hippocampal CA1 area of the *Satb2^flox/flox^* control mice was determined by counting nuclei positive for the SATB2 immunostaining (1:100; Abcam, Cat#ab10563678). To avoid counting the same nuclei, only every 30^th^ ultrathin section cut through the tissue was collected, and the remaining sections discarded. Twenty-five nuclei were identified in the pyramidal neurons layer per image in the DAPI channel only and SATB2-positive cells counted. We confirmed that within the CA1 layer most cells (96%) are pyramidal glutamatergic neurons (data not shown).

For DNs and OLGs, cryosections were washed (3x, 30min total) in PBS, quenched (20 min) in PBS containing 20mM glycine, then permeabilized (15 min) in 0.1% Triton X-100 in PBS (v/v). Cryosections were then incubated (1h, room temperature) in blocking solution (1% BSA (w/v), 0.2% fish-skin gelatin (w/v), 0.05% casein (w/v) and 0.05% Tween-20 (v/v) in PBS). After incubation (overnight, 4°C) with the antibody in blocking solution, the cryosections were washed (3-5x; 1 h) in blocking solution and immunolabeled (1 h, room temperature) with secondary antibodies in blocking solution, followed by 3 (15 min) washes in 0.5% Tween-20 in PBS (v/v). Cryosections were then counterstained with 0.5 µg/ml DAPI in PBS, then rinsed in PBS. Coverslips were mounted in Mowiol® 4-88.

Digital images were acquired with a Leica TCS SP8-STED confocal microscope (Leica Microsystems) using a 63x oil-immersion objective (NA = 1.4) or a 20x oil-immersion objective, using pinhole equivalent to 1 Airy disk. Images were acquired using 405 nm excitation and 420-480 nm emission for DAPI; 488 nm excitation and 505-530 nm emission for TH or GFP; and 555 nm excitation and 560 nm emission using a long-pass filter at 1024x1024 pixel resolution. Images were processed using Fiji (version 2.0.0-rc-69/1.52p), where adjustments included the optimization of the dynamic signal range with contrast stretching.

### Immunofluorescence detection for laser microdissection

For laser microdissection, cryosections on PEN membranes were washed, permeabilized and blocked as for confocal microscopy, and incubated with primary and secondary antibodies as indicated above except for the use of higher concentrations of primary antibodies, as follows: anti-TH (1:50), anti-pan-histone (1:50) or anti-GFP (1:50). Secondary antibodies were used at the same concentration. Cell staining was visualized using a Leica laser microdissection microscope (Leica Microsystems, LMD7000) using a 63x dry objective. Following detection of cellular sections of the cell types of choice containing nuclear slices (nuclear profiles; NPs), individual NPs were laser microdissected from the PEN membrane, and collected into PCR adhesive caps (AdhesiveStrip 8C opaque; Carl Zeiss Microscopy #415190-9161-000). We used multiplexGAM^9^ where three NPs are collected into each adhesive cap, and the presence of NPs in each lid was confirmed with a 5x objective using a 420-480 nm emission filter. Control lids not containing nuclear profiles (water controls) were included for each dataset collection to keep track of contamination and noise amplification of whole genome amplification and library reactions, and can be found in **Supplemental Table 2**.

### Whole genome amplification of nuclear profiles

Whole genome amplification (WGA) was performed using an in-house protocol. Briefly, NPs were lysed directly in the PCR adhesive caps for 4 or 24h at 60°C in 1.2x lysis buffer (30 mM Tris-HCl pH 8.0, 2 mM EDTA pH 8.0, 800 mM Guanidinium-HCl, 5 % (v/v) Tween 20, 0.5 % (v/v) Triton X-100), containing 2.116 units/ml QIAGEN **protease (**Qiagen, Cat#19155). After protease inactivation at 75**°**C for 30 min, the extracted DNA was amplified using random hexamer primers with an adaptor sequence. The pre-amplification step was done using 2x DeepVent mix (2x Thermo polymerase buffer (10x), 400**µ**m dNTPs, 4 mM MgSO_4_ in ultrapure water), **µ**M GAT-7N primers (5′- GTG AGT GAT GGT TGA GGT AGT GTG GAG NNN NNN N) and 2 units/**µ**l DeepVent^®^ (exo-) DNA polymerase (New England Biolabs, M0259L) in the programmable thermal cycler for 11 cycles. Primers that anneal to the general adaptor sequence were then used in a second exponential amplification reaction to increase the amount of product. The exponential amplification was done using 2x DeepVent mix, 10 mM dNTPs, 100 **µ**M GAM-COM primers (5′-GTG AGT GAT GGT TGA GGT AGT GTG GAG) and 2 units/**µ**l DeepVent (exo-) DNA polymerase in the programmable thermal cycler for 26 cycles. For some NPs from DNs (see Supplemental Table 1), WGA was performed using a WGA4 kit (Sigma-Aldrich) using the manufacturer’s instructions.

### GAM library preparation and high-throughput sequencing

Following WGA, the samples were purified using SPRI beads (0.725 or 1.7 ratio of beads per sample volume). The DNA concentration of each purified sample was measured using the Quant-iT® Pico Green dsDNA assay kit (Invitrogen #P7589) according to manufacturer’s instructions. GAM libraries were prepared using the Illumina Nextera XT library preparation kit (Illumina #FC-131-1096) following the manufacturer’s instructions with an 80% reduced volume of reagents. Following library preparation, the DNA was purified using SPRI beads (1.7 ratio of beads per sample volume) and the concentration for each sample was measured using the Quant-iT® PicoGreen dsDNA assay. An equal amount of DNA from each sample was pooled together (up to 196 samples), and the final pool was additionally purified three times using the SPRI beads (1.7 ratio of beads per sample volume). The final pool of libraries was analyzed using DNA High Sensitivity on-chip electrophoresis on an Agilent 2100 Bioanalyzer to confirm removal of primer dimers and estimate the average size and DNA fragment size distribution in the pool. NGS libraries were sequenced on Illumina NextSeq 500 machine, according to manufacturer’s instructions, using single-end 75 bp reads. The number of sequenced reads for each sample can be found in **Supplemental Table 2**.

### GAM data sequence alignment

Sequenced reads from each GAM library were mapped to the mouse genome assembly GRCm38 (Dec. 2011, mm10) with Bowtie2 using default settings^70^. All non-uniquely mapped reads, reads with mapping quality <20 and PCR duplicates were excluded from further analyses.

### GAM data window calling and sample quality control

Positive genomic windows that are present within ultrathin nuclear slices were identified for each GAM library. Briefly, the genome was split into equal-sized windows (50 kb), and number of nucleotides sequenced in each bin was calculated for each GAM sample with bedtools^71^. Next, we determined the percentage of orphan windows (i.e. positive windows that were flanked by two adjacent negative windows) for every percentile of the nucleotide coverage distribution and identified the percentile with the lowest percent of orphan windows for each GAM sample in the dataset. The number of nucleotides that corresponds to the percentile with lowest percent of orphan windows in each sample was used as optimal coverage threshold for window identification in each sample. Windows were called positive if the number of nucleotides sequenced in each bin was greater than the determined optimal threshold.

Each dataset was assessed for quality control by determining the percentage of orphan windows in each sample, number of uniquely mapped reads to the mouse genome, and correlations from cross-well contamination for every sample (**Supplemental Table 2**). Most GAM libraries passed the quality control analyses (86-96% in each dataset; **Extended Data Fig. 1b-d**). To assess the quality of sampling in each GAM dataset, we measured the frequency with which all possible intra-chromosomal pairs of genomic windows are found in the same GAM sample, and found that 98.8 – 99.9% of all mappable pairs of windows were sampled at least once at resolution 50 kb at all genomic distances. Each sample was considered to be of good quality if they had <70% orphan windows, >50,000 uniquely mapped reads, and a cross-well contamination score determined per collection plate of <0.4 (Jaccard index). The number of samples in each cell type passing quality control is summarized in **Table 1**.

### Publicly available GAM datasets from mouse embryonic stem cells (mESC)

For mESCs, GAM datasets were downloaded from 4D Nucleome portal (https://data.4dnucleome.org/). We used 249 x 3NP GAM datasets from mESCs (clone 46C) which were grown at 37°C in a 5% CO_2_ incubator in Glasgow Modified Eagle’s Medium, supplemented with 10% fetal bovine serum, 2 ng/ml LIF and 1 mM 2-mercaptoethanol, on 0.1% gelatin-coated dishes. Cells were passaged every other day. After the last passage 24 h before harvesting, mESCs were re-plated in serum- free ESGRO Complete Clonal Grade medium (Merck, Cat# SF001- B). The list of 4DN sample IDs is provided in **Supplemental Table 6**.

### Visualization of pairwise chromatin contact matrices

To visualize GAM data, contact matrices were calculated using pointwise mutual information (PMI) for all pairs of windows genome-wide. PMI describes the difference between the probability of a pair of genomic windows being found in the same NP given both their joint distribution and their individual distributions across all NPs. PMI was calculated by the following formula, where p(x) and p(y) are the individual distributions of genomic windows x and y, respectively; and p(x,y) are their joint distribution:

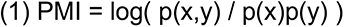

PMI can be bounded between -1 and 1, to produce a normalized PMI (NPMI) value given by the following formula:

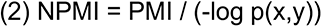

### Insulation score and topological domain boundary calling

TAD calling was performed by calculating insulation scores in NPMI GAM contact matrices at 50 kb resolution, as previously described^3, 9^. The insulation score was computed individually for each cell type and biological replicate, with insulation square sizes ranging from 100 to 1000 kb. TAD boundaries were called using a 500kb-insulation square size and based on local minima of the insulation score. Within each dataset, boundaries that were touching or overlapping by at least 1 nucleotide were merged. Boundaries were further refined to consider only the minimum insulation score within the boundary and 1 window on each side, to produce a 3 bin “minimum insulation score” boundary. In comparisons of boundaries between different datasets, boundaries were considered different when separated by at least one genomic bin, i.e. if they are separated by at least 50 kb. In **Fig. 2c,** we considered the boundary coordinate as the genomic window within a boundary with the lowest insulation value. TAD border coordinates for all cell types can be found in **Supplemental Table 7**. UpSet plots for TAD border overlaps, compartments, and TF motif analyses were generated either with custom Python or R scripts, or using the *UpSetR* tool^72^.

### mESC cell culture for scRNA-seq

Mouse embryonic stem cells from the 46C clone, derived from E14tg2a and expressing GFP under *Sox1* promoter^73^, were a kind gift from D. Henrique (Instituto de Medicina Molecular, Faculdade Medicina Lisboa, Lisbon, Portugal). mESCs were cultured as previously described^19^, i.e. cells were routinely grown at 37°C, 5% (v/v) CO_2_, on gelatine-coated (0.1% v/v) Nunc T25 flasks in GMEM medium (Invitrogen, Cat# 21710025), supplemented with 10% (v/v) Foetal Calf Serum (FCS; BioScience LifeSciences, Cat# 7.01, batch number 110006), 2000 U/ml Leukaemia inhibitory factor (LIF, Millipore, Cat# ESG1107), 0.1 mM beta-mercaptoethanol (Invitrogen, Cat# 31350-010), 2 mM L-glutamine (Invitrogen, Cat# 25030-024), 1 mM sodium pyruvate (Invitrogen, Cat# 11360039), 1% penicillin-streptomycin (Invitrogen, Cat# 15140122), 1% MEM Non- Essential Amino Acids (Invitrogen, Cat# 11140035). Medium was changed every day and cells were split every other day. mESC batches all tested negative for Mycoplasma infection, performed according to the manufacturer’s instructions (AppliChem, Cat#A3744,0020). Before collecting material for scRNA-seq, cells were grown for 48h in serum-free ESGRO Complete Clonal Grade Medium (Merck, Cat# SF001- B) supplemented with 1000 U/ml LIF, on gelatine (Sigma, Cat# G1393-100ml)-coated (0.1% v/v) Nunc 10 cm dishes, with a medium change after 24h.

### Single-cell mRNA library preparation

Two batches (denoted batch A and B, respectively) of single-cell mRNA-seq libraries were prepared according to Fluidigm manual “Using the C1 Single-Cell Auto Prep System to Generate mRNA from Single Cells and Libraries for Sequencing”. Cell suspension was loaded on 10-17 microns C1 Single-Cell Auto Prep IFC (Fluidigm, Cat# 100-5760, kit Cat# 100-6201). After loading, the chip was observed under the microscope to score cells as singlets, doublets, multiplets, debris or other. The chip was then loaded again on Fluidigm C1 and cDNA was synthesised and pre-amplified in the chip using Clontech SMARTer kit (Takara Clontech, Cat# 634833). In batch B, we included Spike-In Mix 1 (1:1000; Life Technologies, Cat# 4456740), as from the Fluidigm manual. Illumina sequencing libraries were prepared with Nextera XT kit (Illumina, Cat# FC- 131-1096) and Nextera Index Kit (Illumina, Cat# FC-131- 1002), as previously described^74^. Libraries from each microfluidic chip (96 cells) were pooled and sequenced on 4 lanes on Illumina HiSeq 2000, 2x100bp paired-end (batch A) or 1 lane on Illumina HiSeq 2000, 2x125bp paired-end (batch B) in the Wellcome Trust Sanger Institute Sequencing Facility (**Supplemental Table 8**).

### ScRNA-seq data processing: mapping and expression estimates

To calculate expression estimates, mRNA-seq reads were mapped with STAR (Spliced Transcripts Alignment to a Reference, v2.4.2a)^75^ and processed with RSEM, using the ‘single-cell-prior’ option (RNA-Seq by Expectation- Maximization, v1.2.25)^76^. The references provided to STAR and RSEM were the gtf annotation from UCSC Known Genes (mm10, version 6) and the associated isoform-gene relationship information from the Known Isoforms table (UCSC), adding information for ERCCs sequences in samples from batch B. Tables were downloaded from the UCSC Table browser (http://genome.ucsc.edu/cgi-bin/hgTables) and, for ERCCs, from ThermoFisher website (www.thermofisher.com/order/catalog/product/4456739). Gene-level expression estimates in “Expected Counts” from RSEM were used for the analysis.

### ScRNA-seq data processing: quality control

Cells scored as doublets, multiplets, or debris during visual inspection of the C1 chip were excluded from the analysis. Datasets were also excluded if any of the following conditions were met: <500,000 reads (calculated using sam-stats from ea-utils.1.1.2-537)^77^; <60% of reads mapped (calculated with sam-stats); <50% reads mapped to mRNA (picard-tools-2.5.0, broadinstitute.github.io/picard/); >15% of reads mapped to chrM (sam-stats); if present, >20% of reads mapped to ERCCs (sam- stats). Correlations between previously published mESCs (clone 46C) mRNA-seq bulk^19^ and the single-cell RNAseq mESC transcriptomes were performed to assess quality of the single cell data. Correlations were performed as previously described^78^. Average single cell expression was highly correlated with bulk RNA-seq (**Extended Data Fig. 2b**).

### ScRNA-seq integration and analysis

To integrate single cell transcriptomes from brain cell types of interest, we selected p21-22 OLGs^16^, p22-32 CA1 PGNs^17^, and p21-26 VTA DNs^18^, based on the cell type and subtype definitions provided in their respective publications. The matrices of counts provided in each publication, along with the single-cell mESC transcriptomes produced which passed QC, were normalized applying the LogNormalize method and scaled using Seurat^79^. The scaled data was used for a PCA analysis, followed by processing through dimensionality reduction using Uniform Manifold Approximation and Projection (UMAP)^80^. Subsequent clustering was performed using the Seurat R package^79^, with default parameters. Single mESC transcriptomes from batch A and B clustered together, and were pooled for further analyses. Genes that could not be mapped to the chosen reference GTF were removed (UCSC; accessed from iGenomes July 17, 2015; https://support.illumina.com/sequencing/sequencing_software/igenome.html.

To generate bigwig tracks for visualization, raw fastq files from each cell type were pooled into one fastq file. Reads were mapped to the mouse genome (mm10) using STAR with default parameters but --outFilterMultimapNmax 10. Bam files were sorted and indexed using Samtools^81^ and normalized (RPKM) bigwigs were generated using Deeptools^82^ bamCoverage. To account for differences in the number of technical replicates in OLG samples, cells were divided into groups by number of runs (1,2 and 6). The median of the reads for the group with the lowest sequencing depth was used as a threshold to normalize the other groups (i.e. the rest of the fastq files were randomly downsampled to that number of reads). The three groups of raw reads were pooled together and processed by applying the same method as for the other cell types. Pseudo-bulk expression was determined using the Regularized Log (R-log) value for each gene (**Extended Data Fig. 2e-f).** In each cell type, only the genes with R-log values ≥2.5 in all pseudo-bulk replicates were considered expressed.

### Differential gene expression analysis

For differential expression analysis for all cell types, pseudo-bulk replicate samples were obtained by randomly partitioning the total number of single cells per dataset into three groups and pooling all UMIs per gene of cells belonging to the same replicate. To determine differentially expressed (DE) genes, all 6 possible pairwise comparisons between samples were performed using DEseq2 with default parameters^83^. Genes classified as DE in at least one comparison were considered for further analysis (adjusted p-value < 0.05; Benjamini-Hochberg multiple testing correction method). A summary table for the DE expression analysis of all cell types can be found in **Supplemental Table 9**. For transcription factor motif analysis, only the DE genes obtained from the comparison between DNs and PGNs were considered for further analysis (**Extended Data Fig. 5c**).

For *Olfr* differential expression analyses, PGNs were classified as positive for *Olfr* expression if at least one gene was detected (≥1 UMI per cell; 164 cells). Two pseudo-replicates were generated for *Olfr*-positive and Olfr-negative cells by dividing the cells in each class in two random groups and pooling all UMIs per gene. DEseq2 was applied to obtain DE genes (adjusted p-value < 0.05; Benjamini-Hochberg method)^83^. To obtain robust results, this process was performed 21 times and only the genes classified as DE in at least 14/21 (67%) of iterations were considered as DE genes for this analysis. A summary table for the robust genes found in the DE expression analysis can be found in **Supplemental Table 10.** The same method was applied for *Vmn* genes (total *Vmn*-expressing cells, 203 cells), *Vmn*-expressing cells that do not express *Olfr* genes (*Vmn* only; 135 cells, 10 iterations only), and for random subsets of cells with the same number of *Olfr*-positive cells (164 cells, 10 iterations only). For *Vmn* only and random subsets, genes classified as DE in at least 7/10 (70%) of iterations were considered as DE genes. Only 12 DE genes were found when comparing total *Vmn*-expressing PGNs to *Vmn-* cells (*Kcnq1ot1*, *Meg3*, *Sngh11*, and *Srrm2* were upregulated; *Aldoa*, *Basp1, Calm2, Lars2, Pfn2, Rtn1, Scg2* and *Sub1* were downregulated; **Extended Data Fig. 9f**). For *Vmn* only cells, only 2 DE genes were found (*Vmn2r29* upregulated; *Lars2* downregulated), and no robust differential genes were detected for the randomly partitioned cells.

### Gene Ontology

Gene ontology (GO) term enrichment analysis was performed using GOElite (version 1.2.4)^84^. In **Fig. 2d**, all genes expressed in at least one cell type, annotated to mm10, were used as the background dataset. In **Fig. 4d**, all genes expressed in PGNs or DNs were used as the background dataset, in **Figs. 5c**, **5d** all unique genes were used, and in **Fig. 5i** all expressed PGN genes were used. Default parameters were used for the GO enrichment: GO terms that were enriched above the background (significant permuted p-values <0.05, 2000 permutations) were pruned to select the terms with the largest Z-score (>1.96) relative to all corresponding child or parent paths in a network of related terms (genes changed >2). GO terms which had a permuted p-value ≥ 0.01, contained fewer than 6 genes per GO term or from the ‘cellular_component’ ontology were not reported in the main figures. A full list of unfiltered GO terms can be found in **Supplemental Table 11**.

### Modeling and in-silico GAM

To reconstruct 3D conformations of the *Nrxn3* locus we employed the Strings & Binders Switch (SBS) polymer model of chromatin^85, 86^. In the SBS, a chromatin region is modelled as a self-avoiding chain of beads, including different binding sites for diffusing, cognate, molecular binders. Binding sites of the same type can be bridged by their cognate binders, thus driving the polymer folding. The optimal SBS polymers for the *Nrxn3* locus in mESCs and DNs were inferred using PRISMR, a machine-learning-based procedure which finds the minimal arrangement of the polymer binding sites best describing input pairwise contact data, such as Hi-C^31^ or GAM^87^. Here, PRISMR was applied to the GAM experimental data by considering the NPMI normalization on a 4.8Mb region around the *Nrxn3* gene (Chr12: 87,600,000- 92,400,000; mm10) at 50kb resolution in mESCs and DNs. The procedure returned optimal SBS polymer chains made of 1440 beads, including 7 different types of binding sites, in both cell types.

Next, to generate thermodynamic ensembles of 3D conformations of the locus, molecular dynamics simulations were run of the optimal polymers, using the freely available LAMMPS software^88^. In those simulations, the system evolves according to Langevin equation, with dynamics parameters derived from classical polymer physics studies^89^. Polymers are first initialized in self-avoiding conformations and then left to evolve in order to reach their equilibrium globular phase^85^. Beads and binders have the same diameter σ =1, expressed in dimensionless units, and experience a hard-core repulsion by use of a truncated Lennard-Jones (LJ) potential. Analogously, attractive interactions are modelled with short-ranged LJ potentials^85^. A range of affinities between beads and cognate binders were sampled in the weak biochemical range, from 3.0K_B_T to 8.0K_B_T (where K_B_ is the Boltzmann constant and T the system temperature). In addition, binders interact unspecifically with the polymer with a lower affinity, sampled from 0K_B_T to 2.7K_B_T. For sake of simplicity, the same affinity strengths were used for all different binding site types. Total binder concentration was taken above the polymer coil-globule transition threshold^85^. For each of the considered cases, ensembles of up to ∼500 distinct equilibrium configurations were derived. Full details about the model and simulations are discussed in Barbieri et al.^85^ and Chiariello *et al*.^90^.

*In-silico* GAM NPMI matrices were obtained from the ensemble of 3D structures by applying the *in-silico* GAM algorithm^10^, here generalized to simulate the GAM protocol with 3NPs per GAM sample and to perform NPMI normalization. Specifically, the same number of slices were used as in the GAM experiments, 249x3NPs for mESCs and 585x3NPs for DNs. Pearson correlation coefficients were used to compare the *in-silico* and experimental NPMI GAM matrices.

Example of single 3D conformations were rendered by a third-order spline of the polymer beads positions, with regions of interest highlighted in different colors. To quantify the size and variability of the 3D structures in mESCs and DNs, the average gyration radius (R_g_) was measured from the selected domains encompassing and surrounding the *Nrxn3* gene, expressed in dimensionless units σ in **Fig. 3e**, **Extended Data Fig. 4d**.

### Melting TAD discovery

To assess gene insulation differences, insulation square values at ten length scales (100 – 1000 kb) were calculated for genes >400 kb in length (n = 479). Cumulative probability distributions of insulation square values were calculated for each dataset, and the brain cells were compared to mESC probability distributions for each gene by computing the maximum distance between the distributions and applying a Kolmogorov-Smirnov test with a custom script in R. P-values were corrected for multiple testing using the Bonferroni method, and -log10 transformed to obtain a ‘TAD melting score’. TAD melting scores for each gene in each comparison can be found in **Supplemental Table 3**. For visualization, empirical cumulative probabilities and insulation score values were smoothed using a Guassian kernel density estimate (adjust = 0.3).

To compare TAD melting scores with transcription across the entire gene, length scaled RPM values were obtained for genes longer than 400 kb. Coordinates of genes were obtained from patch 6 (2017 release) of the Genome Reference Consortium m38 build for improved accuracy. RefSeq assembly accession GCF_000001635.26 was downloaded from NCBI on the 2019-11-19. The number of reads overlapping each gene in each cell type was computed using SAMtools^81^ with the following parameters: *samtools view -b -h -c -F 4*. To obtain Reads Per Million (RPM) in each cell type, the number of overlapping reads was divided by a per million scaling factor (SF) which is specific for the sequencing depth of each cell type (SF = total number of counts / 10^6^). Length-scaled RPM values were obtained per gene with the following formula:

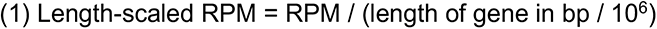

### Determining differential contacts between GAM datasets

To determine significant differences in pairwise contacts between a pair of GAM datasets, genomic windows with low detection, defined as less than 2% of the distribution of all detected genomic windows for each chromosome, were removed from both datasets to be compared. Next, NPMI contact frequencies at each genomic distance of each chromosome were normalized by computing a Z-score transformation, and a differential matrix (D) was derived by subtracting the two Z-score normalized matrices. A 10 Mb distance threshold was applied after computing the normalized matrices. The top significant differential contacts were determined for each dataset by fitting a normal distribution for each chromosome in matrix D and defining the upper and lower 5% of values from the fitted curve.

### Transcription factor binding site analysis

To find transcription factor binding (TF) motifs present within specific contacts, significant differential contacts were determined for DNs and PGNs. For each set of differential contacts, we also constructed three sets of randomly generated contacts, containing the same number of contacts at each genomic distance for each chromosome. Next, accessible regions within the significant contacts were determined using scATAC-seq^37^ or sc low-input chromatin accessibility (liDNase- seq)^38^ for PGNs and DNs, respectively. To account for methodological differences, including lower sequencing depth in DNs, we considered only the PGN scATAC-seq peaks that occurred in >25 cells (of 273 total cells). Motif finding within accessible regions in significant contacts was performed using the Regulatory Genomics Toolbox (https://www.regulatory-genomics.org/motif-analysis/introduction/) with TF motifs (from the HOCOMOCO database, v11)^91^ obtained for TFs expressed in either DNs or PGNs (R-log ≥ 2.5) to determine the percentage of windows containing each TF motif. Next, TF motifs were filtered based on percentage of windows containing the motif (>5%) and differential expression in either PGNs or DNs (-log10(adj. p value) > 3) resulting in 48 TF motifs for feature pair analysis (32 TF motifs from PGNs and 16 from DNs; see **Extended Data Fig. 5d-e**).

For feature pair analysis, we considered the co-occurrence of motif pairs <**i**,**j**> in contacting windows <**w**,**x**>, wherein motif **i** occurs in window **w** and motif **j** occurs in window **x**. The number of contacts containing each pair of selected TF motifs (1128 heterotypic motif pairs and 48 homotypic motif pairs; 1176 total) was calculated, together with the percentage of total significant differential contacts in PGNs and DNs. These metrics were used to calculate an ‘Info Gain’ score and an ‘Enrichment’ score for each TF motif pair. The Info Gain score is a measure of how well a given TF motif pair distinguishes two datasets, in this case PGNs and DNs. Info Gain measures the decrease in total entropy that results from classifying each contact based on the presence of a particular TF motif pair. The total entropy (E_T_) is calculated as a function of the number of contacts in significant windows for PGNs (P_A_) and DNs (D_A_), and the total number of contacts in significant windows, T = P_A_ + D_A_:

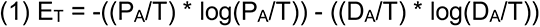

The entropy for a given TF motif pair (E_TF_) was calculated as a function of the number of contacts containing the TF motif pair for PGNs (P_TF_) and DNs (D_TF_), respective to P_A_ and D_A_, given as:

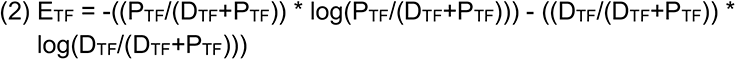

The background entropy (E_B_) for the contacts not containing the TF motif pair for PGNs (P’_TF_ = P_A_-P_TF_) and DNs (D’_TF_ = D_A_-D_TF_) is calculated similarly:

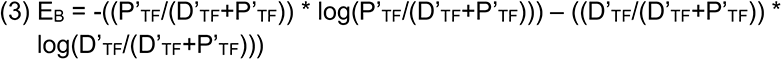

Info Gain was then determined by subtracting the weighted entropies of the TF motif pair and background from the total entropy:

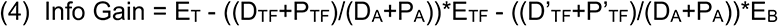

Finally, an Enrichment score was determined by the percentage of contacts in either PGNs or DNs, given by the formula:

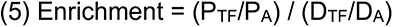

Info gain and Enrichment scores for all TF feature pair interactions, as well as differential expression values (In DNs compared to PGNs) for each TF motif can be found in **Supplemental Table 4.** The top feature pairs were selected based on the top Info Gain scores (10 feature pairs), and highest (PGN enrichment; 5 feature pairs) or lowest (DN enrichment; 5 feature pairs) Enrichment scores. Contact overlaps for top feature pairs were visualized using UpSet plots.

### Network and community detection analysis of transcription factor binding sites in significant differential contacts

To determine the interconnectivity between different TF motifs found in accessible regions of significant differential contacts, the number of contacts for each pair of TF motifs (1176 interactions) was determined. After filtering pairs of TF motifs with <15,000 contacts, a network was built for each cell type with TF motifs as nodes and number of contacts as edges. To detect community membership, the Leiden algorithm was applied using the *leiden* package in R^92, 93^, with a resolution of 1.05 for both PGNs and DNs (**Fig. 4g**, **Supplemental Table 5**).

### Identification of compartments A and B

For compartment analysis, matrices of co-segregation frequency were determined by the ratio of independent occurrence of a single positive window in each sample over the pairwise co-occurrence of pairs of positive windows in a given pair of genomic windows^3^. GAM co-segregation matrices at 250 kb resolution were assigned to either A- or B-compartments, as previously described^3^. Briefly, each chromosome was represented as a matrix of observed interactions O(i,j) between locus i and locus j (co-segregation), and separately for E(i,j) where each pair of genomic windows are the mean number of contacts with the same distance between i and j. A matrix of observed over expected values O/E(i,j) was produced by dividing O by E. A correlation matrix C(i,j) is produced between column i and column j of the O/E matrix. Principal component analysis was performed for the first 3 components on matrix C, before extracting the component with the best correlation to GC content. Loci with PCA eigenvector values with the same sign that correlate best with GC content were called A compartments, while regions with the opposite sign were B compartments. For visualizations and Pearson correlations between datasets, eigenvector values on the same chromosome in compartment A were normalized from 0 to 1, while values on the same chromosome in compartment B were normalized from -1 to 0. Compartments were considered common if they had the same compartment definition within the same genomic bin. Compartment changes between cell types were computed after considering compartments that were common between biological replicates, unless otherwise indicated.

To identify and visualize gene expression differences among genes in changing compartments, K-means clustering was performed on triplicate pseudo- replicates of each cell type using a custom Python script (**Supplemental Fig**. **S8a-b**). The number of clusters were determined using the elbow method, with k-means = 6 for genes in B compartment in mESCs and A in brain cells, and k-means = 5 for A compartment in mESCs and B in brain cells.

### Data availability

Raw fastq sequencing files for all samples from DN, PGN and OLG GAM datasets, together with non-normalized co-segregation matrices, normalized pair- wised chromatin contacts maps and raw GAM segregation tables have been submitted to the GEO repository under accession number GSE94364. Raw fastq sequencing files for mESC GAM datasets are available from 4DN data portal (https://data.4dnucleome.org/). The 4DN sample IDs for all samples used in the study are available in **Supplemental Table 6**. Raw single cell mESC transcriptome data are available from ENA data portal (https://www.ebi.ac.uk/ena/browser/home) The ENA sample IDs for all samples used in the study are available in **Supplemental Table 11**.

