## Supplemental Data Figures & Legends for "Cell-type specialization in the brain is encoded by specific long-range chromatin topologies"

### Extended Data Figure 1

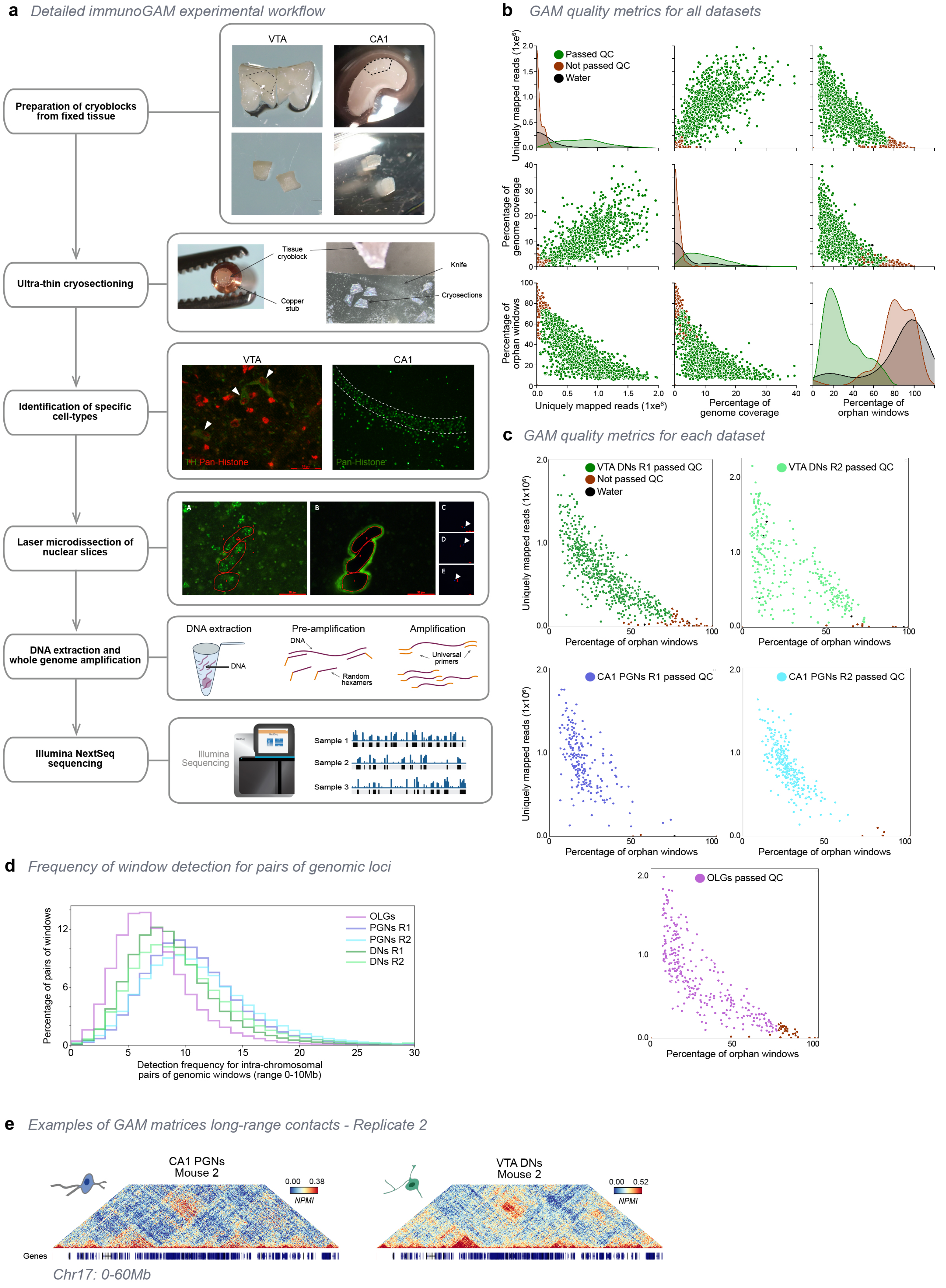

**Extended Data Figure 1. ImmunoGAM experimental pipeline and GAM data quality control.**

**a**, ImmunoGAM experimental pipeline. VTA and CA1 dissections and cryoblock preparations are shown as examples. After fixation, brain tissue is dissected and cryopreserved in sucrose/PBS solution, before sectioning on an ultracryomicrotome (~220nm thick slices; -100ºC). Indirect immunofluorescence using anti-pan-histone antibodies was used to identify cell slices that contained PGN nuclei within the hippocampus, on the laser microdissection microscope, and combined with TH immunolabelling to identify DNs in the midbrain. PGN slices were selected using the morphology of the pyramidal neuron layer. Three nuclear slices were selected and laser microdissected from the tissue to fall into the same PCR lid, as described for multiplex-GAM[9]. Genomic DNA content was then extracted from each sample and amplified using whole-genome amplification, followed by Illumina NextSeq sequencing.

**b**, Correlations between quality control parameters (uniquely mapped reads, genome coverage of positive windows, and percentage of orphan window) for all combined GAM samples collected from brain cell types. Each data point represents a GAM sample. Samples passing QC are shown in green, samples not passing QC in red.

**c**, Correlation between the percentages of uniquely mapped reads and orphan windows per GAM sample shown separately for each dataset produced in this study. Samples not passing QC are shown in red, water control samples (laser microdissected material not containing a nuclear profile) are shown in black.

**d**, Distribution of window detection for pairs of loci at 50-kb resolution (range 0-10 Mb). For all GAM datasets produced in this study, each locus pair is detected an average of 7-10 times.

**e**, Example matrices for CA1 PGN and VTA DN replicate 2 for Chr17: 0-60,000,000 at 50-kb resolution.

#### Extended Data Figure 2

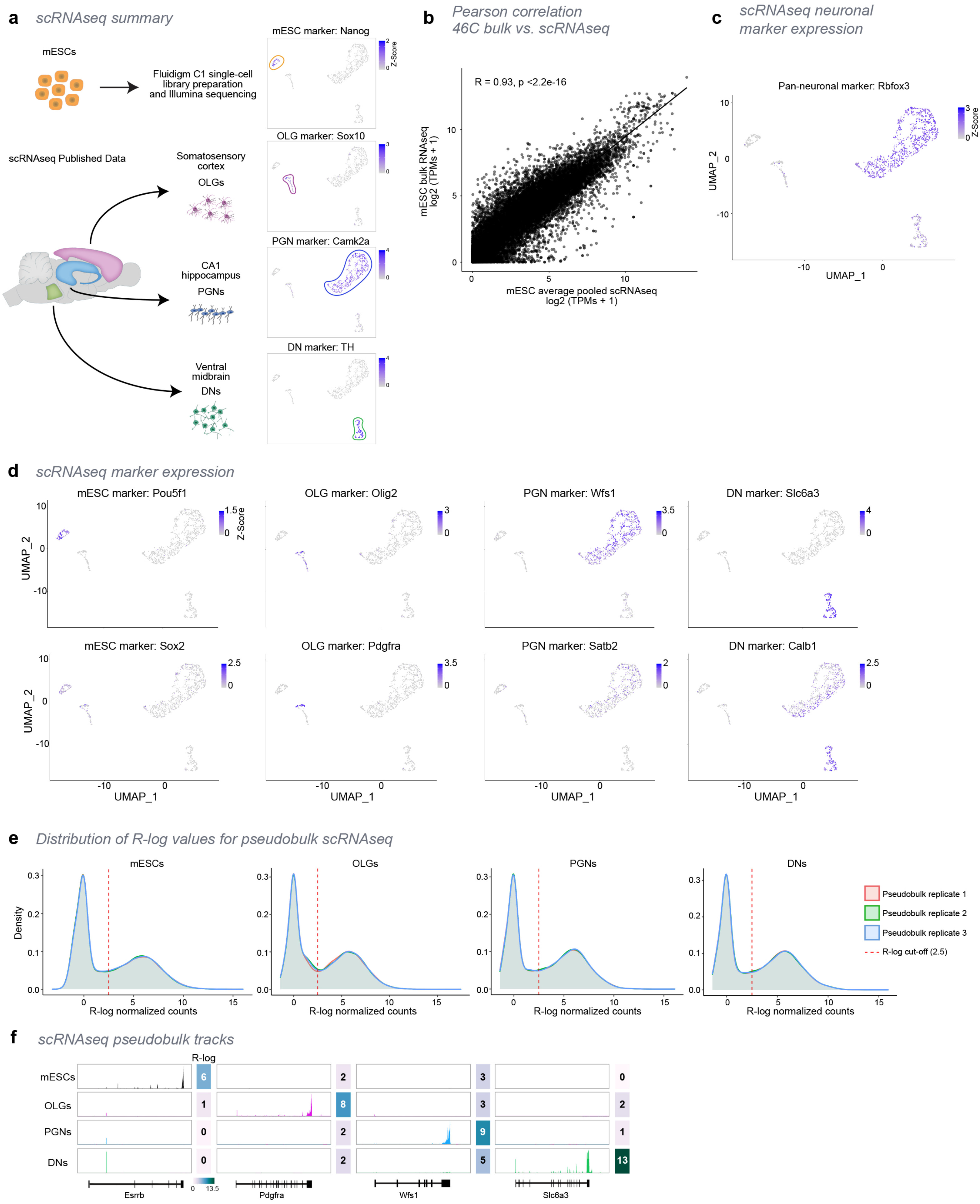

**Extended Data Figure 2. Curation of scRNA-seq data from published datasets in brain cell types and from mESCs.**

**a**, Schematic representation of scRNA-seq datasets used in this study. We generated 98 single-cell transcriptomes from mESCs, and in parallel collected published scRNA-seq datasets from specific cell types in brain regions that were matched with GAM data. Single-cell transcriptomes selected were visualized together by UMAP clustering and colored by expression of each marker gene defined by published data, *Nanog* for mESCs, *Sox10* for OLGs, *Camk2a* for PGNs and *Th* for DNs. Cluster contours are drawn to highlight separation between cell types. All marker genes were found highly expressed in their respective cell types.

**b**, Correlation plot of gene expression in mESCs (clone 46C) between published bulk[19] versus single-cell RNA-seq. Average single-cell expression is highly correlated with bulk RNA-seq.

**c**, Cell-type marker expression, represented by UMAP of single cell transcriptomes for *Rbfox3*, a pan-neuronal marker.

**d**, Additional examples of cell-type marker expression, represented by UMAP of single cell transcriptomes. *Pou5f1* and *Sox2* were used as markers for mESCs, *Olig2* and *Pdgfra* for OLGs, *Wfs1* and *Satb2* for PGNs, and *Slc6a3* and *Calb1* for DNs. All markers are highly expressed in their respective cell types.

**e**, Distribution of regularized log (R-log) values for pseudobulk scRNA-seq datasets. For each cell type, cells were randomly partitioned into 3 pseudobulk replicates before pooling and normalizing reads. The distribution of R-log values is bi-modal for all cell types and pseudobulk replicates. To consider expressed genes for downstream analysis, a 2.5 R-log threshold (dashed red lines) was applied in all datasets. Genes with R-log $\geq$ 2.5 in all three pseudobulk replicates are considered expressed for that cell type.

**f**, Example scRNA-seq pseudobulk tracks for marker genes in each cell type: *Esrrb* for mESCs, *Pdgfra* for OLGs, *Wfs1* for PGNs and *Slc6a3* for DNs. All markers are highly expressed in their respective cell types.

### Extended Data Figure 3

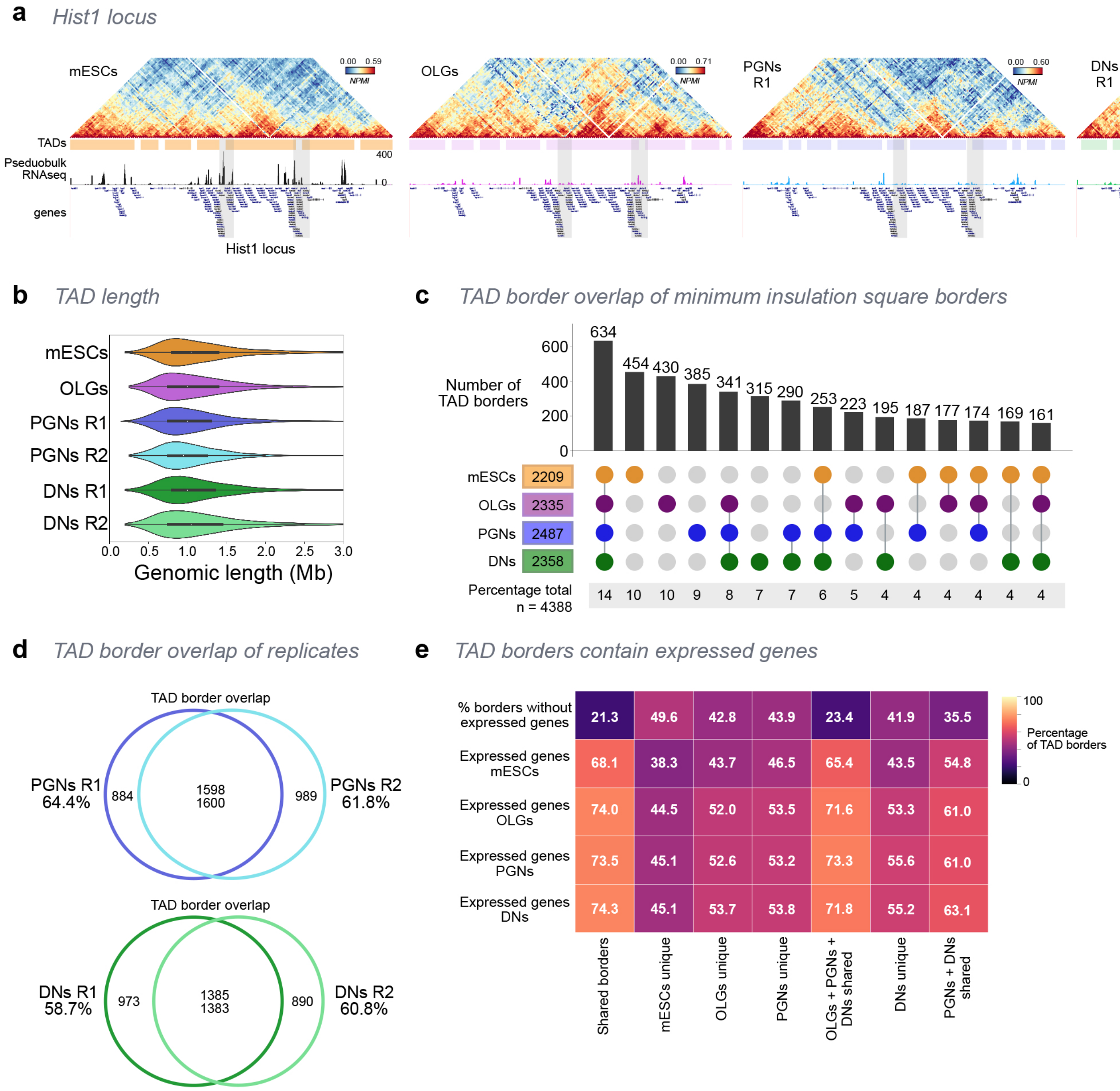


| | Shared borders | mESCs unique | OLGs unique | PGNs unique | OLGs + PGNs + DNs shared | DNs unique | PGNs + DNs shared |
|---|---|---|---|---|---|---|---|
| % borders without expressed genes | 21.3 | 49.6 | 42.8 | 43.9 | 23.4 | 41.9 | 35.5 |
| Expressed genes mESCs | 68.1 | 38.3 | 43.7 | 46.5 | 65.4 | 43.5 | 54.8 |
| Expressed genes OLGs | 74.0 | 44.5 | 52.0 | 53.5 | 71.6 | 53.3 | 61.0 |
| Expressed genes PGNs | 73.5 | 45.1 | 52.6 | 53.2 | 73.3 | 55.6 | 61.0 |
| Expressed genes DNs | 74.3 | 45.1 | 53.7 | 53.8 | 71.8 | 55.2 | 63.1 |

**Extended Data Figure 3. Identification of TADs and TAD boundaries, and differences between cell types.**

**a**, GAM contact matrices show an 8-Mb region surrounding the *Hist1* locus (50-kb resolution; Chr13: 18,000,000-26,000,000). TADs and pseudobulk scRNAseq tracks are indicated for each cell type below matrices. The *Hist1* locus is indicated by the shaded grey areas surrounding a cluster of *Vmn* genes.

**b**, Violin plot shows the distribution of TAD lengths. TAD length was calculated as the distance between two boundary points (defined as lowest insulation score point within a boundary).

**c**, UpSet plot shows TAD boundary overlap for minimum insulation TAD boundaries with at least one cell type containing a TAD border (+1 bin on either side; total 150-kb genomic bins).

**d**, Venn plots show overlap between TAD boundaries in PGN or DN replicates.

**e**, Percentage of TAD borders containing expressed genes (R-log $\geq$ 2.5) in each cell type for the groups shown in (**c**). Higher percentage of borders contain expressed genes in groups with shared borders in two or more cell types. In all groups, brain cells have a higher percentage of borders with expressed genes compared to mESCs.

#### Extended Data Figure 4

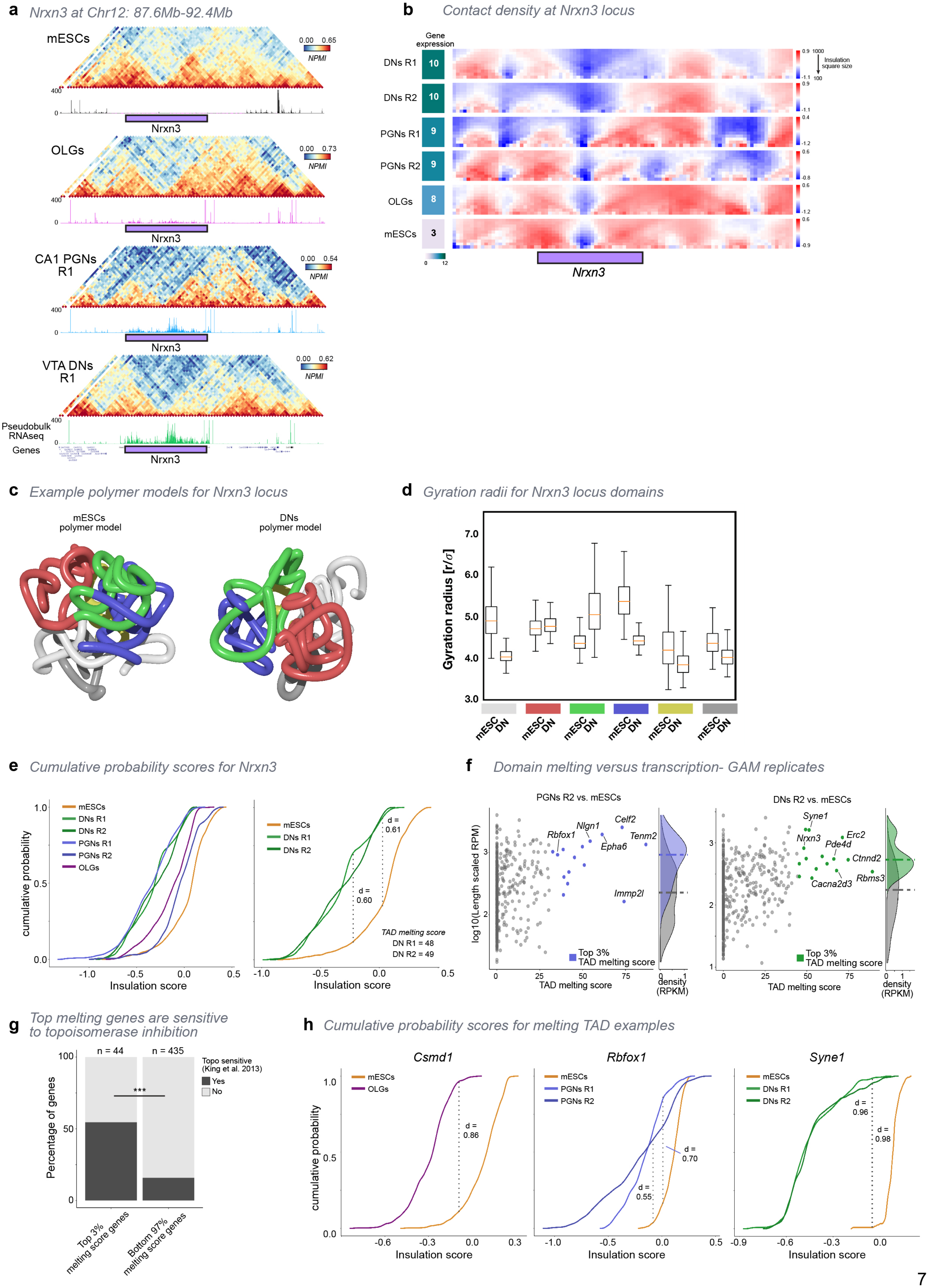

**Extended Data Figure 4. Identification of TAD melting in long expressed genes.**

**a**, GAM contact matrices for a 5Mb region surrounding the *Nrxn3* gene in all cell types (50-kb resolution, chr12: 87,600,000-92,400,000).

**b**, Contact density maps for each cell type and replicate, at the *Nrxn3* locus, calculated using insulation square sizes ranging from 100 - 1000 kb. Contact density is reduced in PGNs and DNs replicate 2 (R2), similar to R1 but occurring in slightly differing regions of the gene.

**c**, Additional examples of polymer models for the *Nrxn3* locus in mESCs and DNs. The *Nrxn3* melted TAD is represented by the green coloured region and is more decondensed in DNs than mESCs. See **Figs**. **3c,d** for location and colouring of the domains.

**d**, Distribution of gyration radii of all domains in polymer models for mESCs and DNs (See **Figs**. **3c,d** for location and colouring of the domains).

**e**, Cumulative probability of insulation square scores ranging from 100 – 1000 kb for *Nrxn3* in all cell types and replicates (*left*). Comparison between DNs R1 and R2 and mESCs, with maximum distance (d) and TAD melting scores (*right*).

**f**, TAD melting scores for each gene (n = 479) in PGNs R2 and DNs R2, compared to mESCs. Genes names with the top 3% of melting scores are coloured in each cell type. Density estimates of length-scaled reads per million (RPM) transcription levels are shown for genes in the top 3% of melting scores (coloured by cell type) compared to genes in the bottom 97% (grey).

**g**, Long genes within the top 3% melting scores (24 of 44 genes) have a higher likelihood of sensitivity to topoisomerase inhibition[28] compared to other genes tested (69 of 435; ***Fisher's exact p-value = $4 \times 10^{-8}$).

**h**, Cumulative probability distributions of insulation scores for genes shown in **Fig. 3h**; *Csmd1* for OLGs, *Rbfox1* for PGNs, and *Syne1* in DNs. All genes were compared to mESCs, with maximum distance (d) indicated for each comparison.

### Extended Data Figure 5

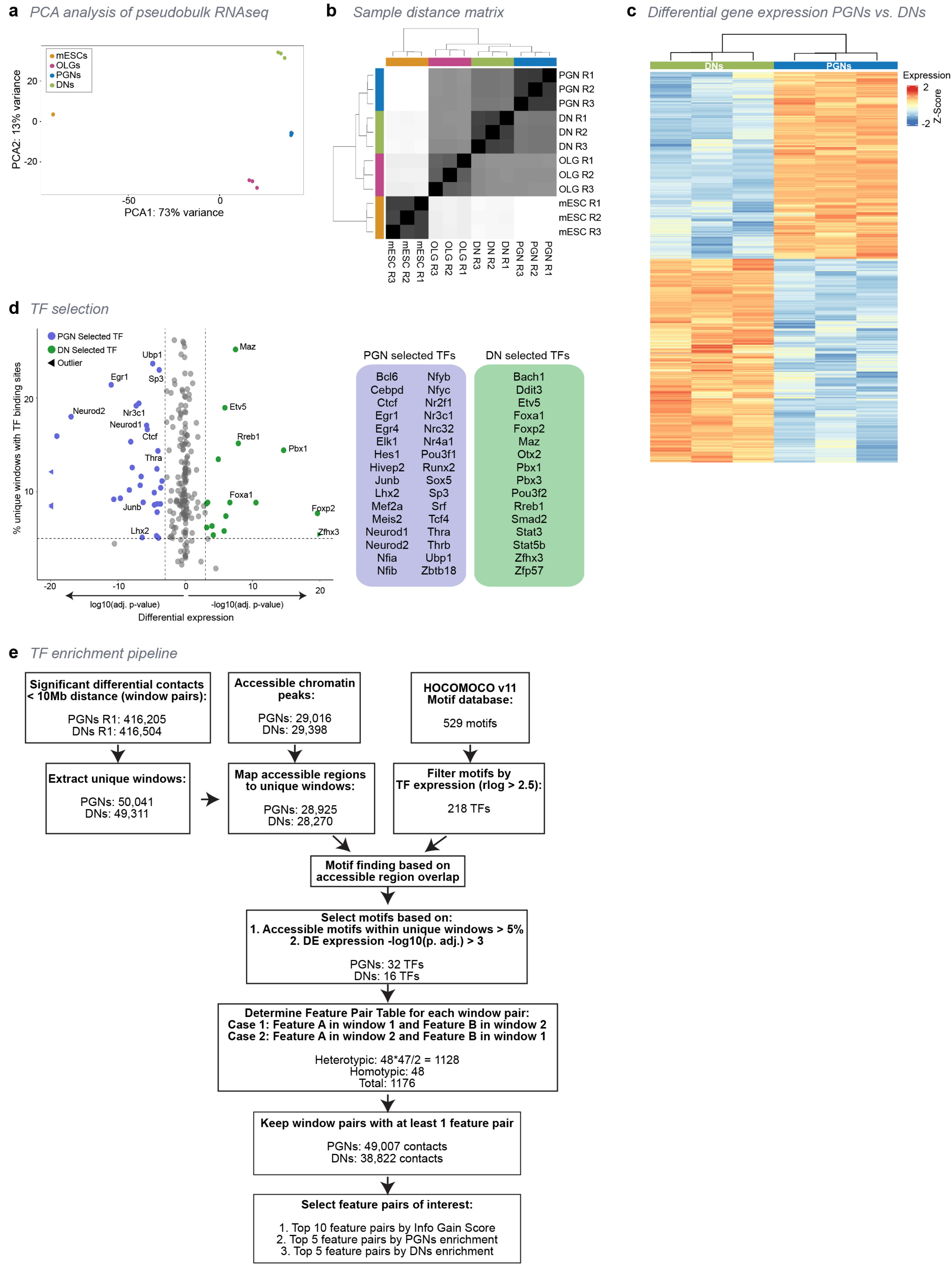

**Extended Data Figure 5. Analysis of transcription factor binding sites and differentially expressed genes in GAM differential contacts.**

**a**, PCA of cell types split into three pseudobulk replicates for differential expression analysis. Pseudobulk replicates clustered separately by cell type.

**b**, Correlation plot of cell type and replicates for differential gene expression analysis. Pseudobulk replicates correlate most highly with one another, followed by brain cell types.

**c**, Heatmap of differentially expressed (DE) genes between PGNs and DNs, clustered by cell type.

**d**, Selection of TF motifs based on percentage of TF motifs in accessible regions within unique windows (> 5%) and differential expression between PGNs (log10(p. adj.) < 3) and DNs (-log10(p. adj.) > 3). PGN-selected TFs (32) are shown in blue, DN-selected TFs (16) are shown in green.

**e**, Full pipeline to determine pairs of transcription factor binding sites in GAM differential contacts. GAM contacts from PGNs and DNs were normalized and compared to produce a differential Z-Score matrix with a 10-Mb distance threshold. The top 5% differential contacts for each dataset were extracted from the differential matrices. Accessible chromatin regions were mapped to the top differential contacts. Next, TF motifs were filtered based on expression in at least one cell type. Accessible region in differential contacts were used to determine the percentage of TF motifs within unique windows. As we were most interested in finding TFs that drive changes if contacts between DNs and PGNs, we chose for further analyses the TF motifs that were found in DN or PGN accessible regions within differential contacts which (1) were present in at least 5% of contacts, and (2) the TFs were differentially expressed between DNs and PGNs (-log10(p.adj.) > 3). The 48 TFs which met the requirements were further investigated to determine the frequency of each motif pair (TF feature pair) in PGN and DN differential contacts. The top-20 TF feature pairs were selected for further analyses based: (a) on Info Gain score (see **Methods** for Info Gain calculation; top 10 feature pairs selected), and (b) on enrichment in either PGNs (top 5 selected) or DNs (top 5 selected).

### Extended Data Figure 6

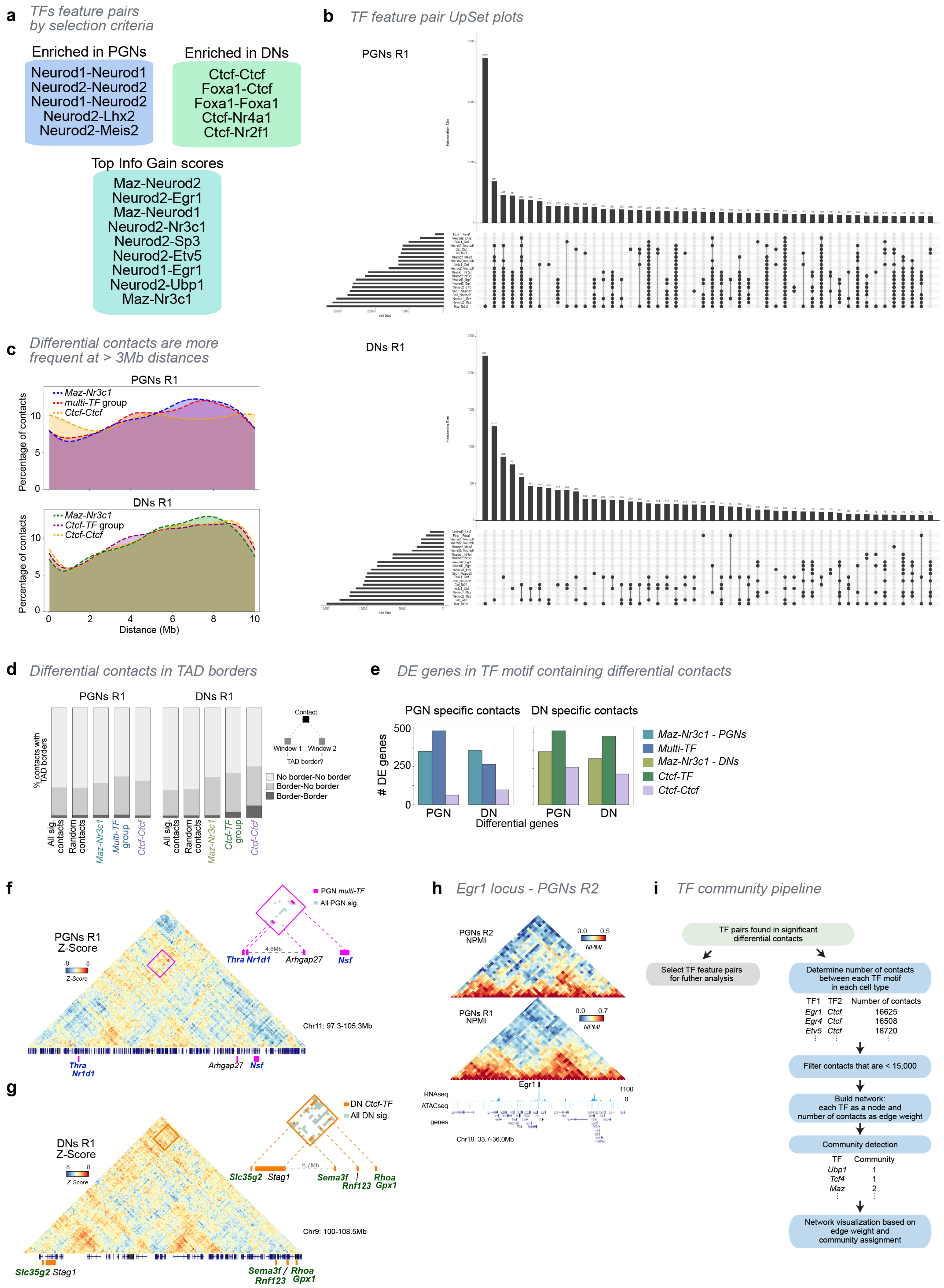

**Extended Data Figure 6. Features of significant differential contacts containing pairs of TF binding sites.**

**a**, TF motif pairs selected by enrichment scores in DNs or PGNs, or by the highest Info Gain scores.

**b**, Overlaps of top 20 TF feature pair contacts for PGN and DN significant differential contacts. The top 50 groups with overlapping TF features are shown for each cell type.

**c**, Percentage of contacts at each genomic distance for significant differential contacts found in TF feature pair groups. Contacts in all groups are enriched at distances > 3 Mb, except for PGN *Ctcf-Ctcf* containing contacts which are found at all distances considered with similar frequency.

**d**, Percentage of contacts (< 2 Mb) that fall within a TAD border in both windows, one window or no windows. For PGNs, most contacts do not overlap with TAD borders, with no differences detected for significant differential contacts found in TF feature pair groups. In DNs, the largest increases are seen in contacts containing the *Ctcf-Ctcf* motif pair, with 47% of the contacts falling in a TAD border.

**e**, Number of PGN or DN differentially expressed (DE) genes found in differential contacts according to sets of TF feature pairs. *Maz-Nr3c1* or *multi-TF* groups of contacts contain several hundred upregulated genes in either cell type. The number of DE genes is lower in *Ctcf-Ctcf* containing contacts.

**f**, Z-Score matrix showing PGN-upregulated genes that form contacts across a ~4.6Mb linear genomic distance (pink box; Chr11: 97,300,000-105,300,000). Inset shows PGN significant differential contacts containing the *multi-TF* group. *Multi-TF* group contacts are shown in pink. Genes highlighted in blue are upregulated in PGNs.

**g**, Z-Score matrix for showing DN-upregulated genes that form contacts across a ~6.7 Mb linear genomic distance (orange box; Chr9: 100,000,000-108,500,000). Inset shows DN significant differential contacts containing the *Ctcf-TF* group. *Ctcf-TF* group contacts are shown in orange. Genes highlighted in green are upregulated in DNs.

**h**, GAM contact matrices showing a 2.3-Mb region surrounding the *Egr1* gene for PGNs R1 and R2 (Chr18: 33,700,000-36,000,000).

**i**, TF motif network and community analysis. After determining the number of contacts for each TF pair, only pairs with > 15,000 contacts were considered. A network was built with each TF as a node and contacts as the edge weight. Community detection was performed using a Leiden algorithm, before visualizing the network.

### Extended Data Figure 7

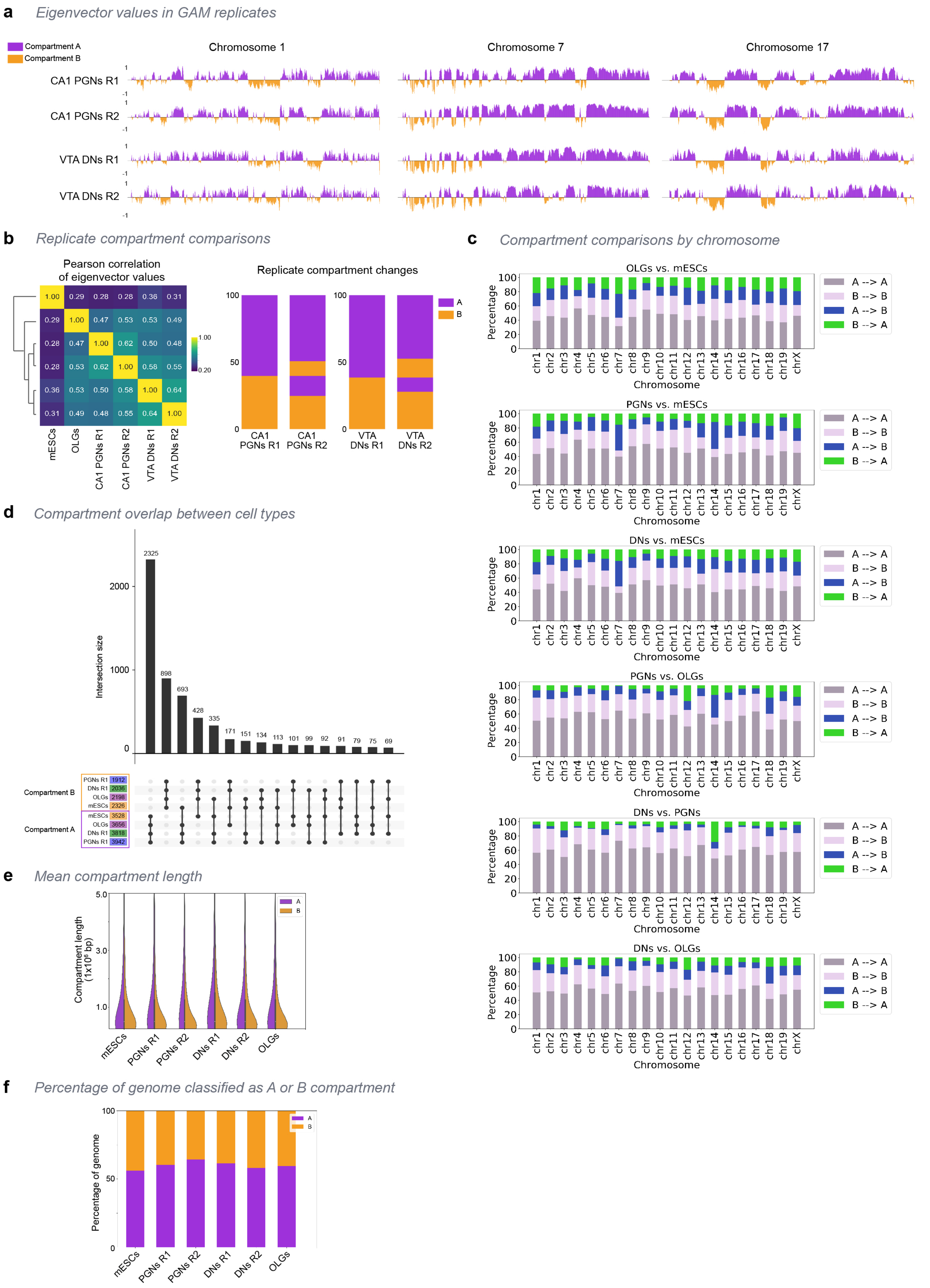

**Extended Data Figure 7. Identification of compartments and differences between cell types.**

**a**, Open and closed chromatin compartments (A and B, respectively) shown for biological replicates of PGNs and DNs. Purple, compartment A; orange, compartment B.

**b**, Correlation of compartments in GAM datasets from different cell types and replicates. Pearson's correlation of eigenvectors shows the largest differences between mESCs and brain cell types. Compartment changes show good overlap between replicates. Purple, compartment A; orange, compartment B.

**c**, Compartment changes for each cell type comparison in each chromosome. Only compartments common to both replicates were used in the comparison. Brain cell types have higher overlap with each other as compared to mESCs. PGNs and DNs had the most overlap for most chromosomes.

**d**, UpSet plot showing all combinations of compartments changes.

**e**, Violin plots of the distribution of compartment lengths show similar lengths between cell types.

**f**, Percentage of the genome covered by A or B compartments in each cell type, showing similar distribution between cell types.

#### Extended Data Figure 8

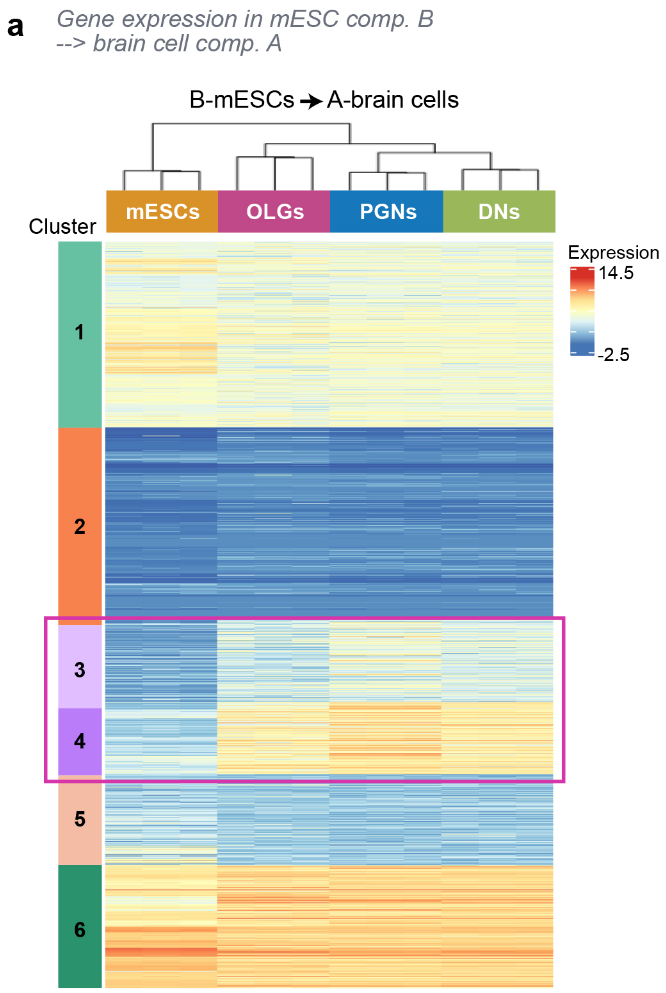


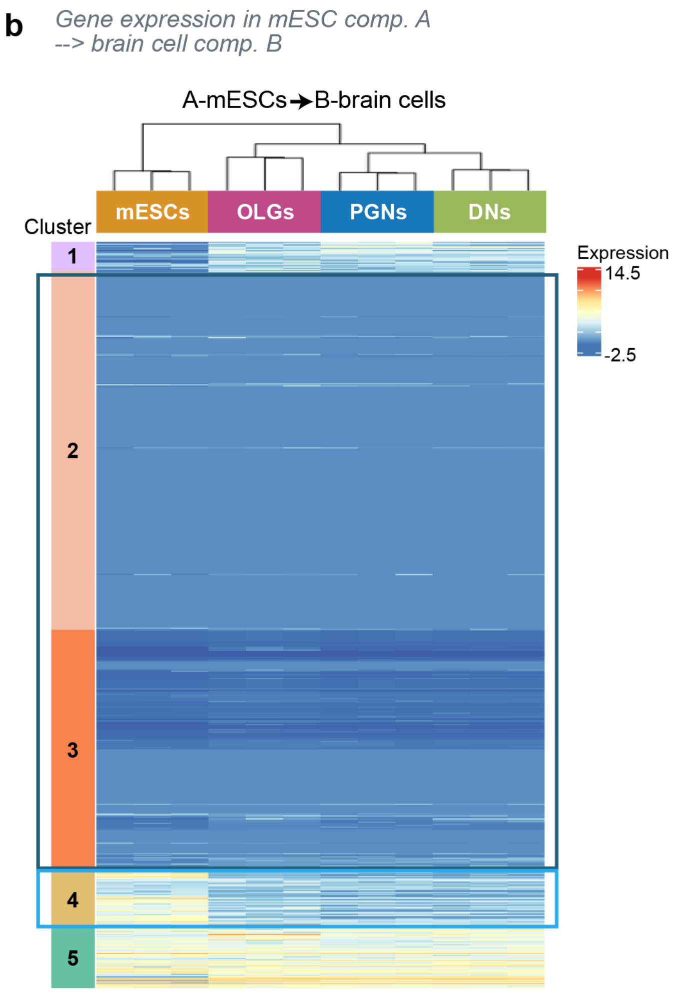

**Extended Data Figure 8. Gene expression and differential gene expression in relation to compartment transitions between mESCs and brain cells.**

**a**, Heatmap of gene expression for genes that change compartments between compartment B in mESCs to compartment A in all brain cells. Clustering of genes by expression shows six distinct clusters where clusters 3 and 4 identify genes that increase their expression between mESC and all brain cell types. Gene ontology (GO) in **Fig. 5c** was done on genes from clusters 3 and 4 combined (pink box).

**b**, Heatmap of gene expression for genes that change from compartment A in mESCs to compartment B in brain cells. Clustering of genes by expression identifies five clusters. Genes in cluster 4 are expressed in mESCs and show lower expression in the brain cell types; they were used for GO analysis presented in **Fig. 5c** (light blue box). Genes in clusters 2 and 3 are not expressed in mESCs nor brain cells; they were combined and used for GO analyses presented in **Fig. 5d** (dark blue box).

### Extended Data Figure 9

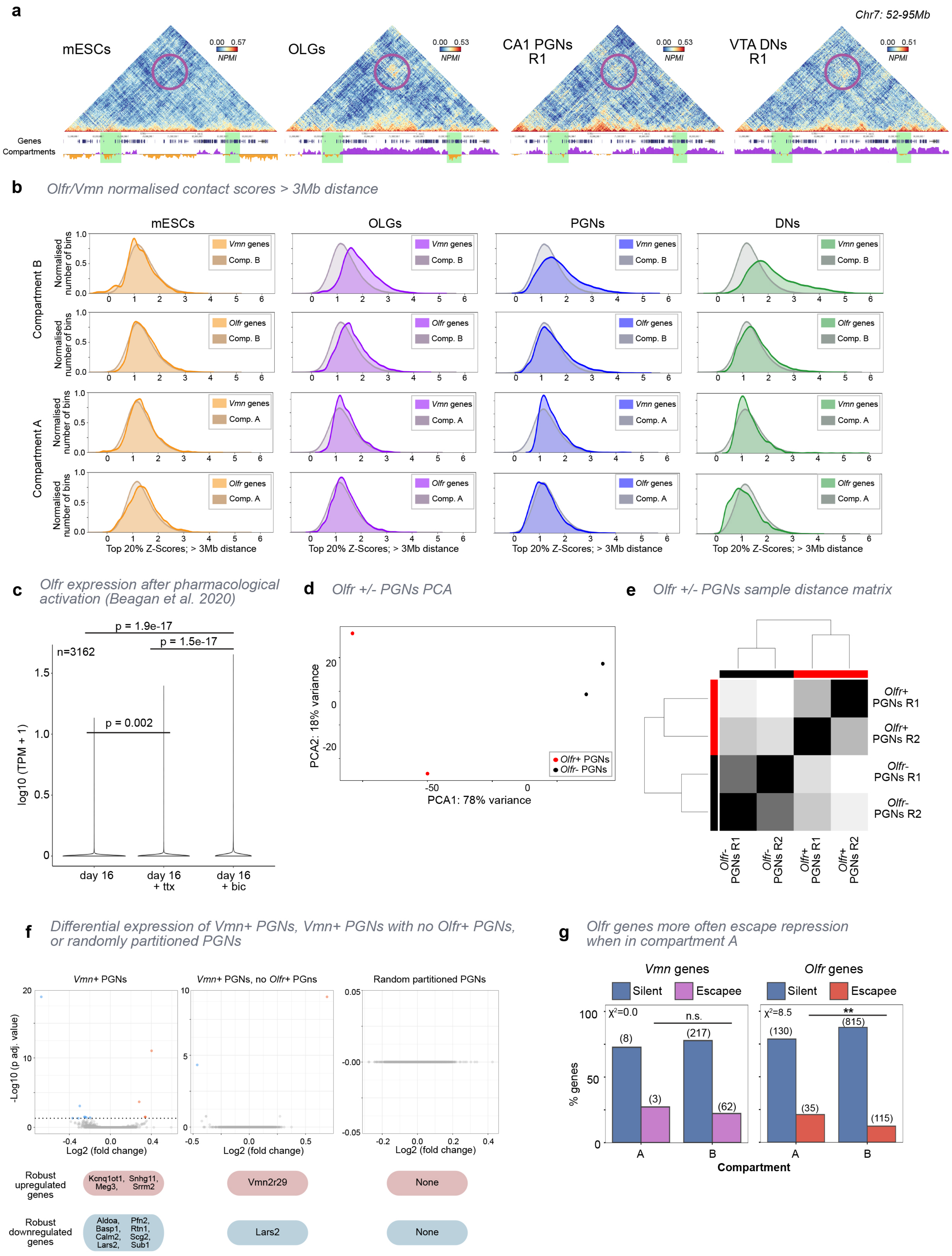

**Extended Data Figure 9. Strong long-range contacts in brain cells regions contain sensory receptor clusters in B compartments.**

**a**, GAM contact matrices show strong interactions that span a 30Mb distance between compartment B regions in OLGs, PGNs and DNs (purple circle), but not mESCs (at 50-kb resolution; Chr7: 52,000,000-95,000,000). Green shaded regions indicate contacts containing *Olfr* and *Vmn* gene clusters.

**b**, Distribution of the top 20% of Z-Score normalized contacts for each genomic window at distances > 3 Mb. For compartment B regions, brain cell types have higher average Z-Scores for compartment windows containing *Vmn* genes, compared to all compartment B windows. Windows containing *Olfr* genes also have higher Z-Score values in OLGs, but the difference is lower in PGNs or DNs. There is no difference for genes found in compartment A for any cell type, or for mESCs in either compartment.

**c**, *Olfr* gene expression is significantly increased after ttx treatment and further increased after bic treatment (significant p-values were determined using a Mann-Whitney-Wilcoxon-test). Transcription levels (log10(TPM)) values for *Olfr* genes were compared between no treatment, or after pharmacological treatment with either tetrodotoxin (ttx), an inhibitor of neuronal firing, or bicuculline (bic), a neuronal activator (RNAseq data from Beagen et al. 2020[7]).

**d**, Exemplar PCA for one iteration of *Olfr*+ and *Olfr*- PGNs split into two replicates for differential expression analysis (see **Methods**). Replicates cluster separately by *Olfr* escapee expression.

**e**, Exemplar correlation plot for one iteration of *Olfr*+ and *Olfr*- PGNs split into two replicates for differential gene expression analysis. Replicates correlate most highly with one another.

**f**, Volcano plot shows one iteration of differentially expressed (DE) genes for PGNs with escapee expression of at least one *Vmn* gene (*Vmn*+ PGNs, *left*), *Vmn*+ PGNs that did not have any *Olfr* gene escape (*middle*), or randomly partitioned PGNs (*right*). Genes were considered to be DE between groups of cells when differential expression had an adjusted p-value < 0.05 in at least 67% of iterations (see **Methods**), and are shown below their respective value in the volcano plot.

**g**, Percentage of *Vmn* or *Olfr* genes that remain silent or escape repression in PGNs for each compartment. For *Vmn*, genes are equally likely to escape repression in either compartment (Chi-squared test, $\chi^2 = 0.0$). *Olfr* genes in compartment A are more likely to escape repression, compared *Olfr* genes in compartment B (Chi-squared test, $\chi^2 = 8.5$; **p = 0.003).
