## Supplementary figures and images for "Cell-type specialization in the brain is encoded by specific long-range chromatin topologies"

### Supplemental Movie 1

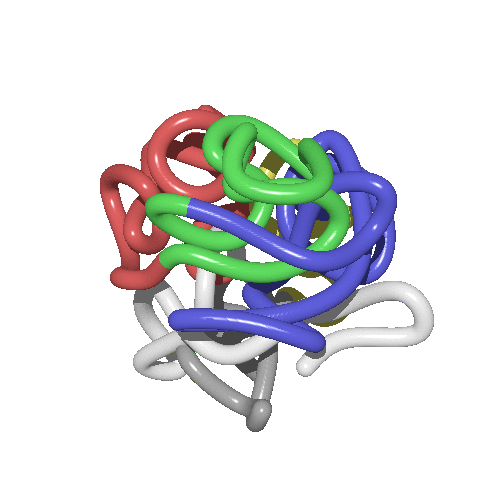

### Supplemental Movie 2

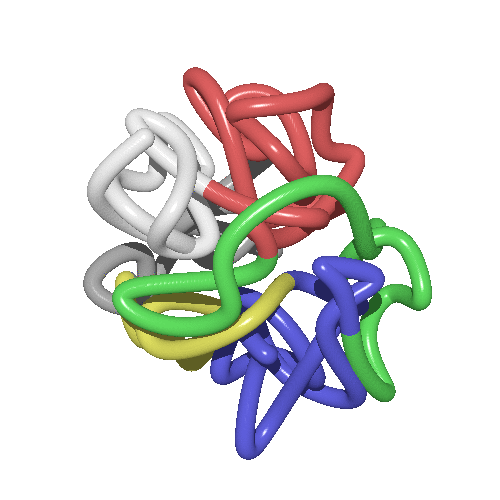
